# Immune signaling induced by plant Toll/interleukin-1 receptor (TIR) domains is thermostable

**DOI:** 10.1101/2024.05.13.592950

**Authors:** Héloïse Demont, Céline Remblière, Laurent Deslandes, Maud Bernoux

## Abstract

Plant disease is a major threat in agriculture and climate change is predicted to intensify it. Above the optimal plant’s growth range, plant immunity and in particular immune responses induced by nucleotide-binding leucine rich repeat receptors (NLRs) are dampened, but the underlying molecular mechanisms remains elusive. NLRs usually contain an N-terminal signaling domain, such as Toll/interleukin-1 receptor (TIR) domain, which is self-sufficient to trigger immune signaling. By using inducible Arabidopsis transgenic lines expressing TIR-containing NLRs (TNLs) or corresponding isolated TIR domains from Arabidopsis RPS4 and flax L6 NLRs, we showed that immune signaling induced downstream of TNL activation is not affected by an elevation of temperature. Conditional activation of TNL- and isolated TIR-mediated immune responses follow the same signaling route at permissive temperature (EDS1/RNLs requirement and activation of the salicylic acid sector). Yet, this signaling pathway is maintained under elevated temperature (30°C) when induced by isolated TIRs, but not full-length TNLs. This work underlines the need to further study how NLRs are impacted by an increase of temperature, which is particularly important to improve the resilience of plant disease resistance in a warming climate.

## Introduction

Climate change intensifies extreme weather conditions such as increased variation of temperature, soil salinity and flooding. Such stresses make plants more vulnerable and threaten ecosystems and agricultural production, which has already been impacted (*1*). Notably, plant disease is currently a major threat for agriculture (*2*), and climate change is predicted to intensify crop losses due to plant pathogens (*3*). The average global temperature is expected to keep rising over the years. Yet, plant immunity is negatively impacted above the optimal temperature plant growth range (*4, 5*).

The plant immune system relies on two major types of immune receptors to fight off pathogens efficiently (*6*). At the cell surface, pathogen-recognition receptors (PRRs) recognize microbial- or damaged-associated molecular patterns (MAMPs/DAMPs). Upon ligand binding, PRRs induce a response called pattern-triggered immunity (PTI). Intracellular receptors from the nucleotide-binding leucine-rich repeats receptors (NLRs) family recognize microbial virulence protein effectors that are secreted inside the cell, which induces effector-triggered immunity (ETI). PTI and ETI are closely interconnected and mutually potentiate each other to mount a robust immune response that includes calcium influxes, Reactive Oxygen Species (ROS) burst, Mitogen-activated protein kinases (MAPK) activation, and a massive transcriptional reprogramming, which sometimes culminates into cell death -the so-called hypersensitive response (HR)-, to locally prevent pathogens progression (*7*). However, this sophisticated immune network can be modulated by temperature changes, by impacting global immune transcriptomic remodeling (*8*), PRR-induced defense responses (*9, 10*) but also NLR-mediated immune outputs, which are often dampened upon an increase of temperature (*11–16*). Given the importance of NLRs in crop breeding programs, the negative impact of temperature elevation on NLR-mediated immunity is a matter of concern in the current context of global warming. However, the molecular mechanisms involved are poorly understood.

NLRs are modular proteins that are commonly composed of three domains. The C-terminal domain has a leucine-rich repeat structure that can be involved in effector sensing. The central nucleotide-binding domain can control the activation state of NLRs, depending on the nucleotide that is bound to it (ADP: ON state, ATP: OFF state), (*17–19*). The N-terminal domain often acts as a signaling unit, which is mainly found as a Toll/interleukin-1 receptor (TIR) or a coiled-coil (CC) domain (*20–23*). When TIR- and CC-containing NLRs (termed TNLs and CNLs, respectively) sense, directly or indirectly, the presence of pathogen effectors, they are qualified as sensor NLRs. A small subclade of NLRs act downstream of sensor NLRs to transduce immune signaling and are therefore called helper NLRs (*24*). Helper NLRs that display a RPW8-like N-terminal CCR domain (RNLs) play a central role in the plant immune system as they are required for immune signaling induced by multiple sensor NLRs and PRRs (*25–27*)(Collier 2011, Saile 2020, Pruitt et al. 2021).

In the recent years, major discoveries have shed light on NLRs structural information and TIR enzymatic functions (*28*). Upon effector recognition, activation of Arabidopsis CNL ZAR1 and wheat CNL Sr35 leads to the formation of a pentameric structure called a resistosome. CNL oligomerization drives close proximity of the CC domain of each protomer, creating a funnel-like structure that function as Ca^2+^ permeable channel at the plasma membrane, which is necessary for immune signaling activation (*29–31*). Similarly, TNLs such as the Arabidopsis RPP1 and tobacco ROQ1 also form resistosomes upon effector recognition, but with a tetrameric stoichiometry. TNL oligomerization promotes TIR enzymatic NADase activity, which in turn activates the ENHANCED DISEASE SUSCEPTIBLITY 1 (EDS1) family and RNLs (*32–35*). In Arabidopsis, EDS1 forms mutually exclusive heterodimers with the other two members of the family, PHYTOALEXIN DEFICIENT 4 (PAD4) or SENESCENCE-ASSOCIATED GENE101 (SAG101) (*36*).

NAD^+^-derived small molecules specifically bind to either EDS1-PAD4 and EDS1-SAG101 complex, which in turn selectively associate with and activate different RNLs subgroups, ACTIVATED DISEASE RESISTANCE 1 (ADR1s) or N REQUIREMENT GENE 1 (NRG1s), respectively (*37–39*). Upon activation, RNLs form oligomeric complexes that are targeted to membranes to form cation-permeable channels (*40, 41*). Although RNLs function redundantly, ADR1s seem to predominantly activate transcriptional reprogramming, including genes involved in the salicylic acid (SA) biosynthesis pathway to induce basal immunity, while NRG1s favor the activation of cell death (*26, 42*).

Interestingly, TIR domains seem to display dual enzymatic function. The nature of the catalytic site can differ by using NAD^+^ as substrate or by hydrolyzing dsRNA/dsDNA to generate various NAD^+^-derived small molecules or 2′,3′-cAMP/cGMP, respectively (*38, 39, 43*). Both enzymatic activities are required to activate immune signaling, although it is not known how 2′,3′-cAMP/cGMP contribute to signaling activation. SA plays a crucial role in basal immunity and NLR-mediated immune responses. Notably, SA accumulation is promoted by EDS1 and PAD4 activity (*44, 45*). However, accumulating evidence feature SA as a potential Achille’s heel of plant immunity under elevated temperatures. At 30°C, the transcript level of genes encoding central regulators of the SA biosynthesis pathway, such as transcription factor CBP60g, biosynthetic enzyme isochorismate synthase 1 ICS1 but also EDS1 and PAD4, is dramatically reduced or abolished in plants challenged with virulent pathogens (*9, 46*). NLR-mediated resistance is also dampened under elevated temperature (above 28°C) (*5*). However, it is not clear how the heat sensitivity of the SA sector may impact NLR activity or *vice versa*. An increase of temperature seems to affect NLRs subcellular localization, which might disrupt their function (*14, 47*). A thermotolerant variant of the Arabidopsis TNL SNC1 maintains its subcellular localization and can propagate PAD4-dependent immune signaling, including SA marker PR1, at elevated temperature, suggesting that increased temperature might alter receptor functions upstream of induced signaling (*14*). However, the underlying mechanisms are not clearly understood.

As signaling units, TIR domains isolated from TNLs can induce autoimmune responses when overexpressed *in planta* in absence of pathogens (*20, 22, 23, 48–50*) and hence, can be used as a tool to study signaling events downstream of the activation of TNLs. In order to investigate the impact of temperature on TNL function and signaling, we used inducible Arabidopsis transgenic lines expressing TIR domains isolated from different thermosensitive TNLs. By comparing lines expressing isolated TIRs and corresponding full-length TNLs at permissive (21°C) and non-permissive (30°C) temperature, we showed that signaling induced by isolated TIRs is maintained under elevated temperature, and this includes the main known actors involved in TNL signaling, such as EDS1, the SA sector and RNLs. Altogether, our data suggest that sensor NLRs are impacted by environmental factors such as temperature, whereas downstream signaling is more resilient.

## Results

### Cell death induced by isolated TIR domains but not full-length TNLs is maintained under elevated temperature

When overexpressed *in planta*, TIR domains isolated from TNLs are self-sufficient to induce an immune response (*20, 22, 51*). Hence, they represent a useful tool to investigate TNL signaling events independently of effector recognition and NLR activation, by overcoming constraints that can be met with full-length receptors intermolecular interactions and conformational changes. To investigate and compare the impact of temperature on full-length TNL- or TIR-induced signaling, we monitored cell death, a proxy for NLR-induced signaling, upon activation of either autoimmune full-length TNLs or corresponding isolated TIRs, under permissive and non-permissive temperature growth conditions. Under permissive temperature (18-22°C), overexpression of the full-length Arabidopsis TNL RPS4 and the flax TNL L6MHV autoimmune variant (which contains an aspartate to valine substitution in the MHD motif, located in the NB domain) leads to autoimmune responses and cell death in transgenic Arabidopsis lines (*11, 52, 53*). Hence, we used inducible Arabidopsis transgenic lines expressing RPS4 and L6MHV, or their corresponding isolated TIR domains, RPS4^TIR^ and L6^TIR^ ((*53*), Table S1). All gene constructs are under the control of a Dexamethasone (Dex)-inducible promoter in order to conditionally activate TNL- or isolated TIR-mediated signaling, and monitor subsequent cell death. They also harbor a C-terminal FSBP (3xFlag/streptavidin-binding peptide), or 3xmyc tag fusion, to follow protein accumulation. At permissive temperature (21°C), Arabidopsis seedlings carrying *pDex:L6MHV*, *pDex:L6^TIR^* and *pDex:RPS4^TIR^* constructs induced autoimmune responses from 3 to 4 days upon transfer on Dex-containing media (+Dex) to reach seedling death after 7 days (Bernoux et al. 2023, Figure 1A). In contrast, seedlings transferred to non-inducing media (-Dex) showed no phenotype and their development were similar to wild-type Col-0 seedlings (Figure 1A). Similar autoimmune responses were observed in at least one independent transgenic line carrying each construct, after Dex treatment (Bernoux 2023, Figure S1A,B).

**Figure 1.**
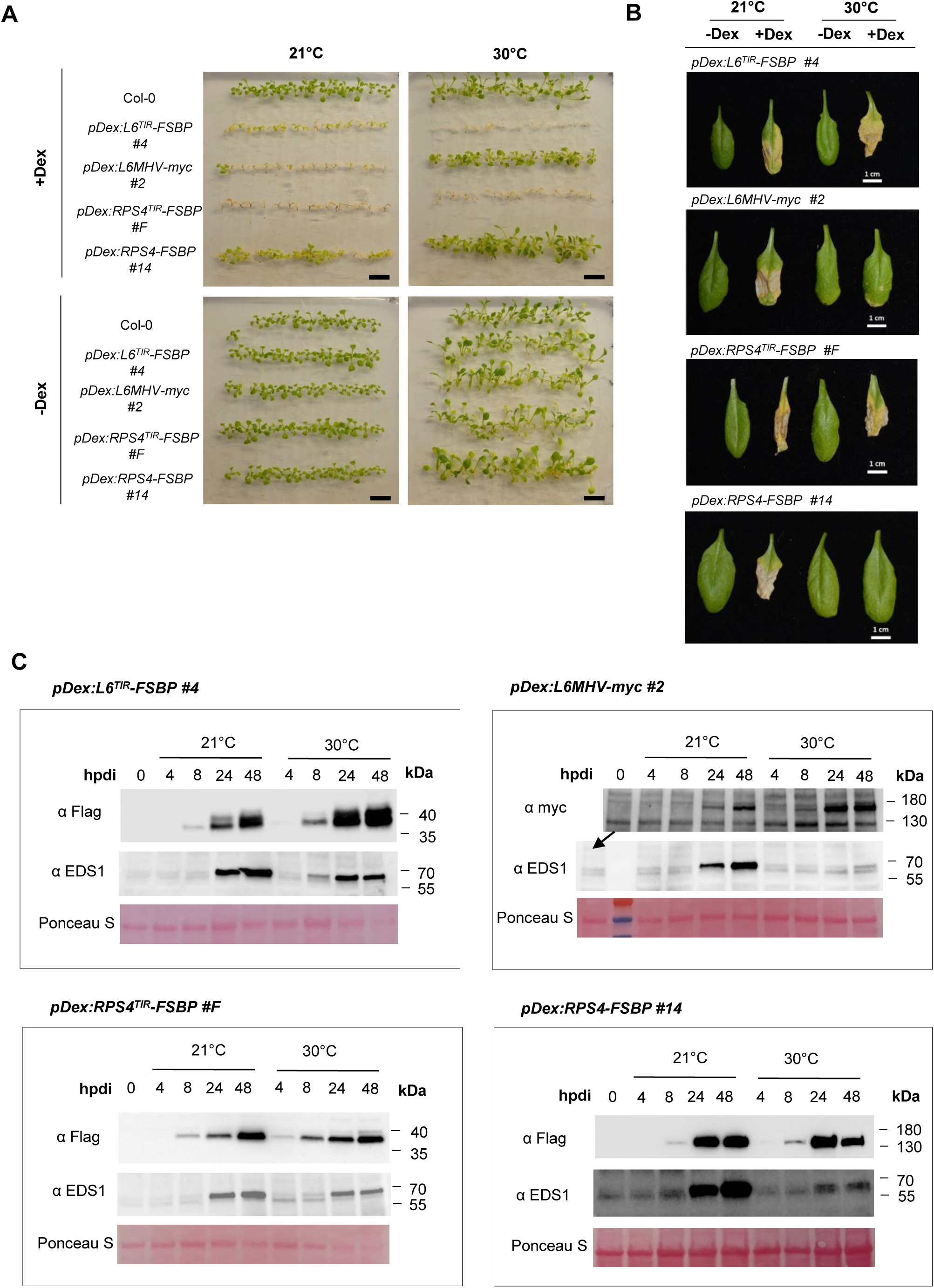
Effect of elevated temperature on cell death induced by TNLs or isolated TIRs in Arabidopsis. (A-B) Cell death phenotype observed on seedlings (A) or adult plant leaves (B) of transgenic inducible lines carrying *pDex:L6^TIR^-FSBP* (line #4), *pDex:L6MHV-myc* (line #2), *pDex:RPS4^TIR^-FSBP* (line #F) and *pDex:RPS4-FSBP* (line #14) after Dex induction. (A) Ten day-old seedling were transferred to 10µM Dex-containing (+Dex) MS media or non-inducing media (DMSO-containing, -Dex) and moved to 21°C or 30°C climatic chambers. Wild-type Col-0 was used as a negative control. Photos were taken seven days after Dex induction. Black scale bars represent 1,5 cm. (B) Leaves of 4 week-old plants were infiltrated with 20 µM Dex (+Dex) or DMSO (-Dex) solutions and moved to 21°C or 30°C climatic chambers. Photos were taken seven days after Dex induction. (C) Immunoblot analysis of L6^TIR^-FSBP, L6MHV-myc, RPS4^TIR^-FSBP, RPS4-FSBP and native EDS1 proteins over a 48h time-course post Dex induction (hpdi) of ten day-old seedlings at 21°C or 30°C, using anti-flag, anti-myc and anti-EDS1 antibodies detection. Total protein load is indicated by red Ponceau staining. Equal amount of protein extract were loaded for each timepoint to detect transgenic protein construct and EDS1.

Seedling cell death was less clear in transgenic lines carrying *pDex:RPS4*. Out of two independent lines (lines #14 and #11), line #14 showed clear autoimmune responses upon seedlings Dex induction, yet did not reach cell death (Figure 1A, Figure S1C). Line #11 did not show any visible symptoms upon Dex induction in seedlings, which may be due to low level of protein accumulation in these conditions, even at higher Dex concentration (100µM) (Figure S1C,D). Cell death was clearly visible on leaves of four weeks-old plants of all Dex-inducible lines, 2 to 3 days after infiltration with a 20µM Dex solution (Figure 1B, Figure S2A), including lines carrying *pDex:RPS4*, even though protein accumulation was still very low for line #11 (Figure S2B). These results demonstrate that cell death signaling induced by L6^TIR^, RPS4^TIR^ and L6MHV can be monitored in both seedling and adult plants, whereas cell death can only be reached in adult plants grown in soil when induced by overexpression of RPS4 at 21°C. Consequently, these data indicate that all generated constructs are competent for activation of immune responses under permissive temperature.

As many NLRs, including the RRS1-R/RPS4 NLR pair, are compromised above 28°C (*11, 12*), we compared the response of our inducible transgenic seedlings at 21°C and 30°C upon Dex induction. When seedlings carrying *pDex:L6MHV* or *pDex:RPS4* were transferred at 30°C after Dex induction, immune responses were strongly reduced and only displayed slight stunting phenotype, in comparison to induced seedlings that remained at 21°C (Figure 1A, Figure S1B-C), supporting that both L6 and RPS4-mediated immunity are compromised at elevated temperature in our assay. In contrast, seedlings expressing corresponding TIR domains alone (L6^TIR^ and RPS4 ^TIR^) showed cell death symptoms that were similar or even stronger to what was observed at 21°C (Figure 1A). Immunoblot analyses showed that L6^TIR^, RPS4 ^TIR^ and corresponding full-length proteins L6MHV and RPS4 can be detected from 8 hours post Dex induction (hpdi) and that protein accumulation increases progressively over the time monitored (24 and 48hpdi) (Figure 1C). Protein accumulation kinetics upon Dex induction were unchanged or stronger at 30°C compared to 21°C for all constructs (Figure 1C), demonstrating that the inhibition of cell death phenotype in seedlings carrying *pDex:L6MHV* and *pDex:RPS4* at 30°C is not due to a lack of protein accumulation. Similar phenotypes were observed in adult plant leaves infiltrated with Dex at 21°C or 30°C. While leaves of lines carrying *pDex:L6^TIR^* and *pDex:RPS4^TIR^* showed clear cell death over the infiltration area at both temperatures, lines carrying *pDex:L6MHV* or *pDex:RPS4* showed no symptoms at 30°C, despite protein accumulation at 24hpdi (Figure 1B, Figure S2B). We obtained similar results with independent lines carrying *pDex:L6MHV* or *pDex:RPS4* (Figure S2A, B). Inducible transgenic lines expressing a TIR domain isolated from another TNL from Arabidopsis (SNC1), also showed autoimmune symptoms that were similar or stronger at 30°C compared to 21°C (Figure S3), whereas autoimmune phenotype mediated by a mutation in full-length *SNC1 (snc1-1)* is *reverted at 28°C* (*14, 54*). NLRs activity is tightly controlled, notably via intra-domain interactions, which could be sensitive to temperature changes. Given their simpler architecture, isolated TIRs are not subjected to such negative controls, which could explain their heat stable activity. In order to test this hypothesis, we generated inducible transgenic lines to express naturally occurring TIR-containing truncated TNLs that were previously described to trigger an immune response when overexpressed (*55*), such as TN2 (TIR-NB with no LRR domain) and RBA1 (TIR-only protein with a C-terminal unknown domain). Interestingly, induction of TN2 or RBA1 induced clear autoimmune responses at 30° C when expressed in seedlings or adult plant leaves (Figure S4). Even though the function of canonical TNLs is affected by an elevation of temperature, our results demonstrate that signaling induced by TIR domains is thermostable.

### Enhanced expression of *EDS1* and genes involved in the salicylic acid (SA) sector is maintained under elevated temperature when induced by isolated TIRs but not full-length TNLs

To decipher and compare the impact of temperature on TNL- and isolated TIR-induced signaling, we monitored the expression of marker genes involved in TNL signaling in our inducible transgenic lines expressing full-length TNLs or isolated TIRs at permissive and elevated temperature.

EDS1 is a major node of plant immunity and most TNLs including RPS4, SNC1 and L6 depend on EDS1 to activate immune signaling ((*36, 53, 56*). RPS4^TIR^ and L6^TIR^ -induced cell death phenotype is abrogated in Arabidopsis *eds1-2* mutant background, suggesting that isolated TIR-induced immune responses depend on EDS1 and follow similar signaling routes as sensor TNLs in Arabidopsis ((*50, 53*), Figure S5). By using anti-EDS1 antibodies, we monitored EDS1 protein accumulation in transgenic Arabidopsis seedlings upon Dex induction of full-length TNLs L6MHV and RPS4 or their corresponding isolated TIR domains. Interestingly, at 21°C, native EDS1 protein accumulation was dramatically higher at 24hpdi in all lines, in comparison to non-induced seedlings (0hpdi), and this correlated with the accumulation of inducible protein constructs (full-length L6MHV and RPS4 or isolated TIR domains) (Figure 1C). Interestingly, at 30°C, EDS1 increased accumulation was maintained upon L6^TIR^ and RPS4^TIR^ induction in contrast to lines expressing L6MHV and RPS4, despite similar or even higher protein accumulation at 30°C compared to 21°C (Figure 1C). To decipher whether lack of increased EDS1 protein accumulation at 30°C upon induction of L6MHV and RPS4 was due to transcriptional or translational inhibition, we analyzed by RT-qPCR the effect of elevated temperature on *EDS1* expression at 24hpdi in seedlings carrying full-length *pDex:L6MHV* or *pDex:RPS4*, and *pDex:L6^TIR^* or *pDex:RPS4^TIR^* at 21°C and 30°C. Consistent with the immunoblot data (Figure 1C), *EDS1* expression was upregulated at 21°C in all transgenic lines in comparison to wild-type seedlings, which is in line with previous studies showing that *EDS1* expression is induced in lines constitutively overexpressing RPS4 (Heidrich et al. 2013). However, at 30°C, *EDS1* increased expression was abolished or significantly reduced in seedlings expressing full-length TNLs L6MHV and RPS4, while it was maintained in those expressing L6^TIR^ and RPS4^TIR^ (Figure 2A). These results demonstrate that *EDS1* gene expression and protein accumulation are induced during both TNL- and TIR-only-mediated early immune responses, but that an increase of temperature prevents *EDS1* induced expression mediated by TNLs but not by isolated TIRs. These results further support that temperature elevation disrupts the function and/or properties of full-length TNL receptors rather than the induced downstream signaling.

**Figure 2.**
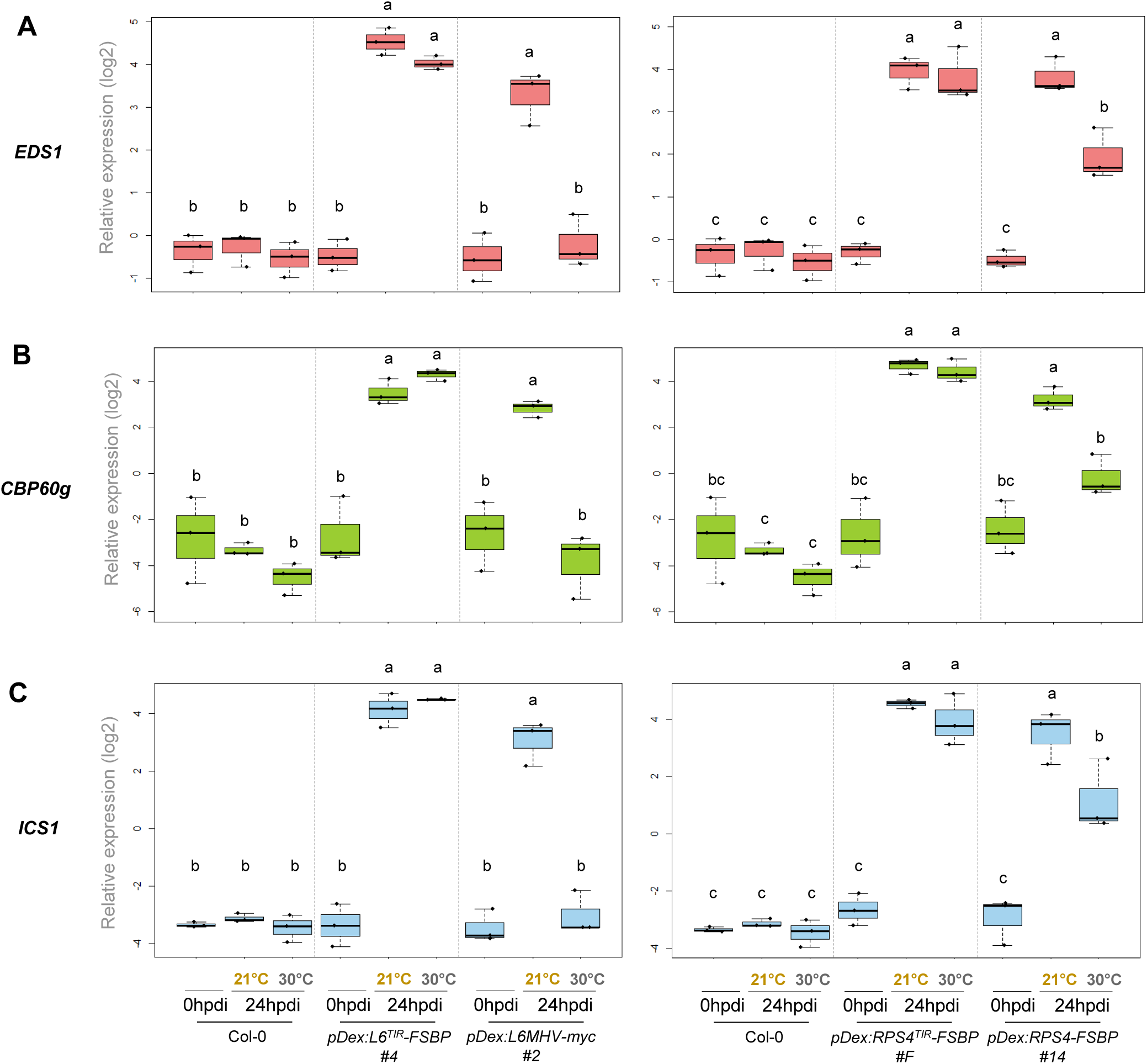
Effect of elevated temperature on the expression of genes involved in the SA pathway induced by TNLs or isolated TIRs. (A-C) RT-qPCR analysis of *EDS1* (A), *CBP60g* (B) and *ICS1* (C) gene expression level in wild-type (Col-0) or transgenic Arabidopsis seedlings carrying *pDex:L6^TIR^-FSBP* (line #4), *pDexL6MHV-myc* (line #2) and *pDex:RPS4^TIR^*-FSBP (line #F) or *pDex:RPS4-FSBP* (line #14) at 21 or 30°C. Seedlings were collected at time 0 and 24 hours after transfer on 10µM Dex-containing medium (hpdi: hours post Dex induction). Gene expression was normalized relatively to the expression of two housekeeping genes (*At1G13320* and *At5G15710)*. Box plots represent values obtained from three independent biological replicates which are represented by black dots. Statistical differences were assessed with two-ways variance analysis (ANOVA) followed by a Tukey’s HSD multiple comparison test. Data points with different letters indicate significant differences (P < 0.05).

EDS1 is a regulator of SA biosynthesis, which is crucial for plants to mount a robust defense (*44, 45*). A recent study highlighted the vulnerability of the SA sector during temperature stress (*46*). Expression of genes encoding major components of the SA biosynthetic pathway, such as *CBP60g* and *ICS1*, is dramatically reduced at 28°C in Arabidopsis plants challenged with a virulent *Pseudomonas syringae* bacterial strain or treated with the SA analog BTH, compared to 21°C, leading to decreased disease resistance (*46*). These findings prompted us to test whether the expression of *CBP60g* and *ICS1* was also impacted by an increase of temperature in our Dex-inducible seedling assay upon TNL or TIR-only activation. RT-qPCR analyses revealed that the expression of both *CBP60g* and *ICS1* was upregulated at 21°C in all transgenic lines expressing full-length TNLs or isolated TIRs, in comparison to wild-type Col-0 seedlings (Figure 2B,C). Strikingly, increased expression of *CBP60g* and *ICS1* was maintained at 30°C in plants expressing L6^TIR^ and RPS4^TIR^, whereas it was reduced to basal levels as in wild-type plants in lines expressing full-length L6MHV or significantly reduced in lines expressing full-length RPS4, which is in line with *EDS1* expression profile (Figure 2). These results show that both TNLs and isolated TIR domains induce the expression of genes involved in SA biosynthesis but in contrast to full-length NLRs, this expression is maintained at elevated temperature when induced by isolated TIR domains. Altogether, our results further support that elevated temperature affects NLRs function but not downstream signaling pathways.

### RNLs are required for signaling induced by both isolated TIR domains and full-length TNLs

TNLs require RNLs to induce immune responses (*57*). Arabidopsis bares five RNLs that are grouped into two subfamilies, ADR1s and NRG1s. ADR1 subfamily is composed of three members, ADR1, ADR1-L1 and ADR1-L2 (*58*), while NRG1 subfamily consists of two members, NRG1.1 and NRG1.2 (*42, 59*). In Arabidopsis, ADR1s and NRG1s function downstream of effector-activated NLRs in an unequally redundant manner, with some functional specificities (*26, 42*). For example, ADR1s mainly contribute to disease resistance induced by the NLR pair RRS1/RPS4, whereas NRG1s seem to translate effector-mediated cell death (*26*). Autoimmune phenotype induced by the *suppressor of npr1-1, constitutive 1 (snc1)* mutant depends on ADR1s, although NRG1s contribute to this phenotype to a lesser extent (*42*). To test whether thermotolerant signaling induced by isolated TIR domains also requires helper NLRs, as for full-length thermosensitive TNLs, we introduced Dex-inducible RPS4^TIR^, SNC1^TIR^, L6^TIR^ and L6MHV constructs into RNL subgroup mutants (*adr1 triple (adr1, adr1-L1, adr1-L2), nrg1 double (nrg1.1, nrg1.2)* and the *helperless* mutant lacking the five RNLs (Saile 2020)). We next monitored cell death symptoms in seedlings from at least two independent transgenic lines for each construct in each genetic background upon Dex induction (Figure 3A, Figures S6-9). In lines carrying *pDex: RPS4^TIR^*, seedlings showed strong cell death symptoms in wild-type and *nrg1 double* mutant background, seven days after Dex induction (Figure 3A, Figure S6A). In the *adr1 triple* mutant, autoimmune phenotype was clear but attenuated compared to wild-type or *nrg1 double* mutant backgrounds, while these symptoms were fully abolished in the *helperless* mutant (Figure 3A, Figure S6A). Immunoblot analyses revealed that RPS4^TIR^ generally accumulated at slightly lower levels in *adr1 triple* and *helperless* mutants compared to as in wild-type or *nrg1 double* mutant (Figure 3B). Because TIR-induced autoimmune responses can be dose-dependent, we performed this seedling cell death assay with increased Dex concentration to increase RPS4^TIR^ protein accumulation in *adr1 triple* and *helperless* transgenic lines. At 50µM Dex, RPS4^TIR^ protein levels in *adr1 triple* and *helperless* mutants were comparable to levels observed in seedlings induced at 10µM Dex in *nrg1 double* mutants (Figure S6B). Consequently, seedlings showed a stronger autoimmune phenotype leading to cell death compared to seedlings induced with 10µM Dex in the *adr1 triple* mutant background (line #332-6)(Figure 3A, Figure S6A), suggesting a dose-dependent effect of RPS4^TIR^ -induced cell death. Overall, these results indicate that both NRG1 and ADR1 families contribute to RPS4^TIR^-mediated immunity.

**Figure 3.**
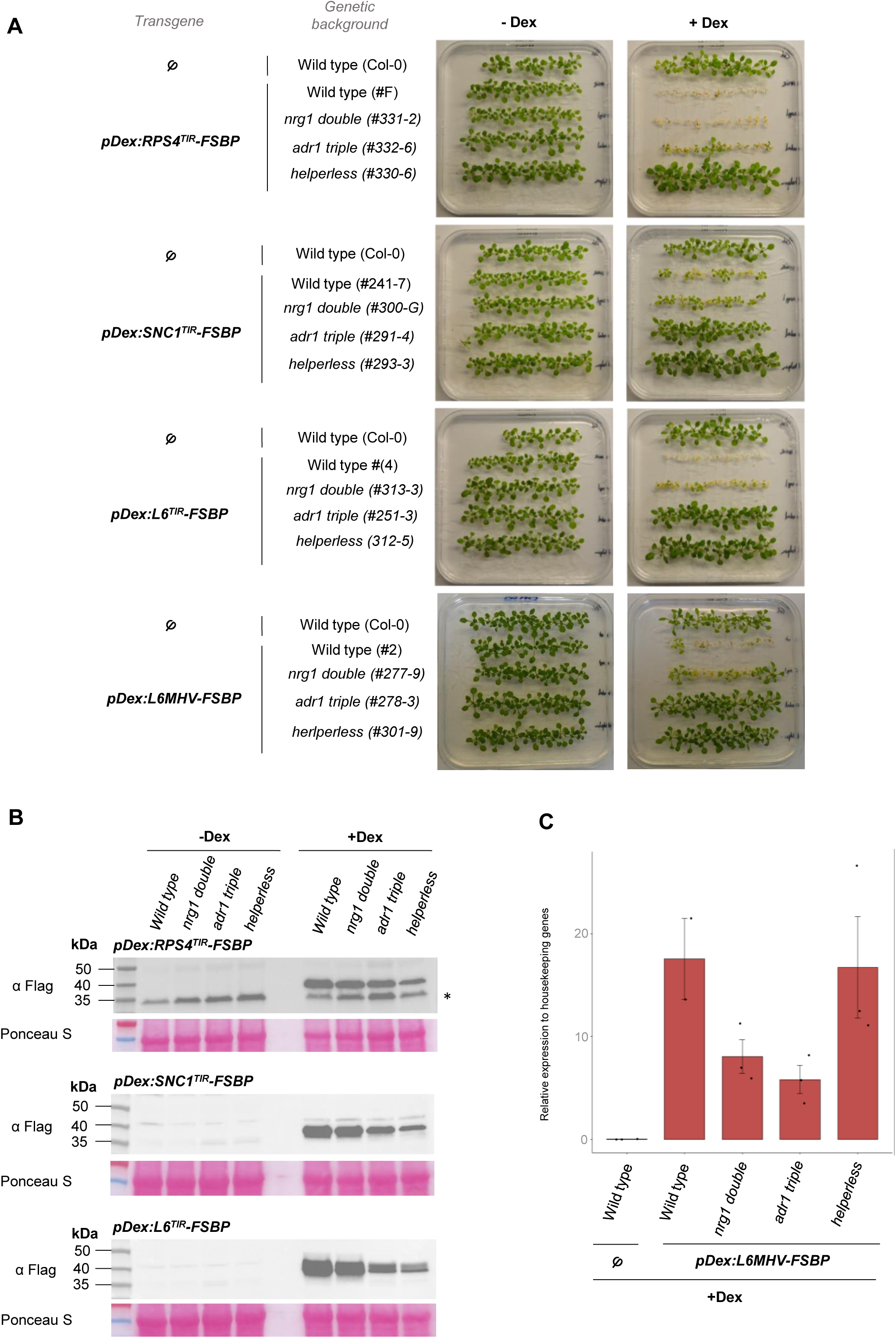
RNLs genetic requirements for signaling induced by TNLs or isolated TIRs. (A) Cell death phenotype observed on seedlings of transgenic inducible lines carrying *pDex:RPS4^TIR^-FSBP*, *pDex:SNC1^TIR^-FSBP*, *pDex:L6^TIR^-FSBP*, or *pDex:L6MHV-FSBP*, in wild-type (wt) or in *RNLs* mutant backgrounds (*nrg1 double*, *adr1 triple* or *helperless*), after Dex induction. Ten day-old seedling were transferred to 10µM Dex-containing (+Dex) MS media or non-inducing media (DMSO-containing, - Dex). Untransformed wild-type Col-0 was used as a negative control. Photos were taken seven days after transfer. (B) Immunoblot analysis of RPS4^TIR^-FSBP, SNC1^TIR^-FSBP and L6^TIR^-FSBP in wild-type or in *RNLs* mutant backgrounds (*nrg1 double*, *adr1 triple* or *helperless*), 24 hours post Dex induction using anti-flag antibodies detection. Total protein load is indicated by red Ponceau staining. Equal amount of protein extract was loaded for each sample. Background noise is indicated by an asterisk. (C) RT-qPCR analysis of *L6MHV-FSBP* transgene expression in wild-type or *RNLs* mutant backgrounds. Untransformed wild-type Col-0 was used as a negative control. *L6MHV-FSBP* expression was normalized relatively to the expression of two housekeeping genes (*At1G13320* and *At5G15710)*. Bar plots represent means +/-SEM obtained from two or three independent biological replicates which are represented by black dots.

In lines carrying *pDex:SNC1^TIR^*, *pDex:L6^TIR^*or *pDex:L6MHV*, seedlings showed autoimmune phenotype that were similar or slightly attenuated in *nrg1 double* compared to wild-type background 7 days upon Dex induction (Figure 3A). At the same time point, no visible or very mild symptoms were visible in the *adr1 triple* mutant background while transgenic seedlings in the *helperless* mutant showed normal development, similarly to non-transgenic wild-type seedlings (Figure 3). Similar trends were observed in at least two independent transgenic lines for each construct and each mutant background (Figures S7-9). Although we attempted to select transgenic lines with similar levels of TIR protein accumulation, we again generally observed a lower level of TIR protein accumulation in *adr1 triple* and *helperless* mutants, compared to *nrg1 double* and wild-type backgrounds, especially for L6^TIR^ (Figure 3B, Figures S7,8). To boost L6^TIR^ protein production, we increased Dex concentration in our induction assay for lines carrying *pDex:L6^TIR^ in adr1 triple* and *helperless* backgrounds. At 50µM Dex concentration, L6^TIR^ protein accumulated at more comparable levels in *adr1 triple (line #251-3)* and *helperless (line #312-5)* mutants relative to *pDex:L6^TIR^* carrying lines in wild-type and *nrg1 double* mutant induced at 10µM Dex. Yet, seedlings from lines in *adr1 triple* and the *helperless* mutant backgrounds still showed no autoimmune phenotype 7 days after 50µM Dex induction, demonstrating that the lack of cell death phenotype in these genetic backgrounds is not due to lower TIR protein accumulation (Figure S8). In lines carrying *pDex:L6MHV*, Dex-induced expression of *L6MHV* transcripts was analyzed by RT-qPCR because protein accumulation was too low for immunodetection. We also observed some differences for *L6MHV* transcript levels in the different mutant backgrounds, with lower expression of *L6MHV* in *adr1 triple* and *nrg1 double* compared to wild-type and *helperless* backgrounds (Figure 3C, Figure S9). Yet, induction of *pDex:L6MHV* led to strong stunting phenotype in the *nrg1 double* mutant, which phenotype was comparable to seedlings carrying *pDex:L6MHV* in wild-type background. In contrast, no autoimmune phenotype was observed in lines carrying *pDex:L6MHV* in *adr1 triple* and *helperless* mutants, as for lines expressing L6^TIR^ and SNC1^TIR^ (Figure 3A).

Taken together, these results show that L6^TIR^, SNC1^TIR^, RPS4^TIR^ and L6MHV depend on both ADR1 and NRG1 RNL subgroups to induce signaling, but lack of NRG1 family has a lesser or no impact compared to the ADR1 family on autoimmune and cell death symptoms induced by SNC1^TIR^, L6^TIR^ and L6MHV. Most importantly, our data show that thermotolerant isolated L6^TIR^ and thermosensitive L6MHV behave similarly in the different mutant backgrounds, suggesting they share the same genetic requirements, such as RNLs, to induce signaling.

### Differential temperature sensitivity of ADR1s-mediated signaling

RNLs function downstream to sensor TNLs when activated by effectors (*26*). We showed here that the ADR1 family mainly contribute to signaling induced by thermosensitive autoimmune TNL L6MHV as well as thermotolerant L6^TIR^ (Figure 3), suggesting that ADR1s must be resilient to an elevation of temperature, unlike sensor TNLs. To test this hypothesis, we looked at the autoimmune responses induced by seedlings carrying *pDex:L6^TIR^*in *nrg1 double* mutant background, which carry wild-type *ADR1s* and show clear cell death at 21°C, under elevated temperature conditions. Interestingly, seven days post-induction, seedlings carrying *pDex:L6^TIR^* displayed clear autoimmune phenotype at 21°C and 30°C in both wild-type and *nrg1 double* mutant background, but not in *adr1 triple* mutant background, demonstrating that ADR1s-dependent *L6^TIR^*-mediated cell death is still effective at 30°C (Figure 4A).

**Figure 4.**
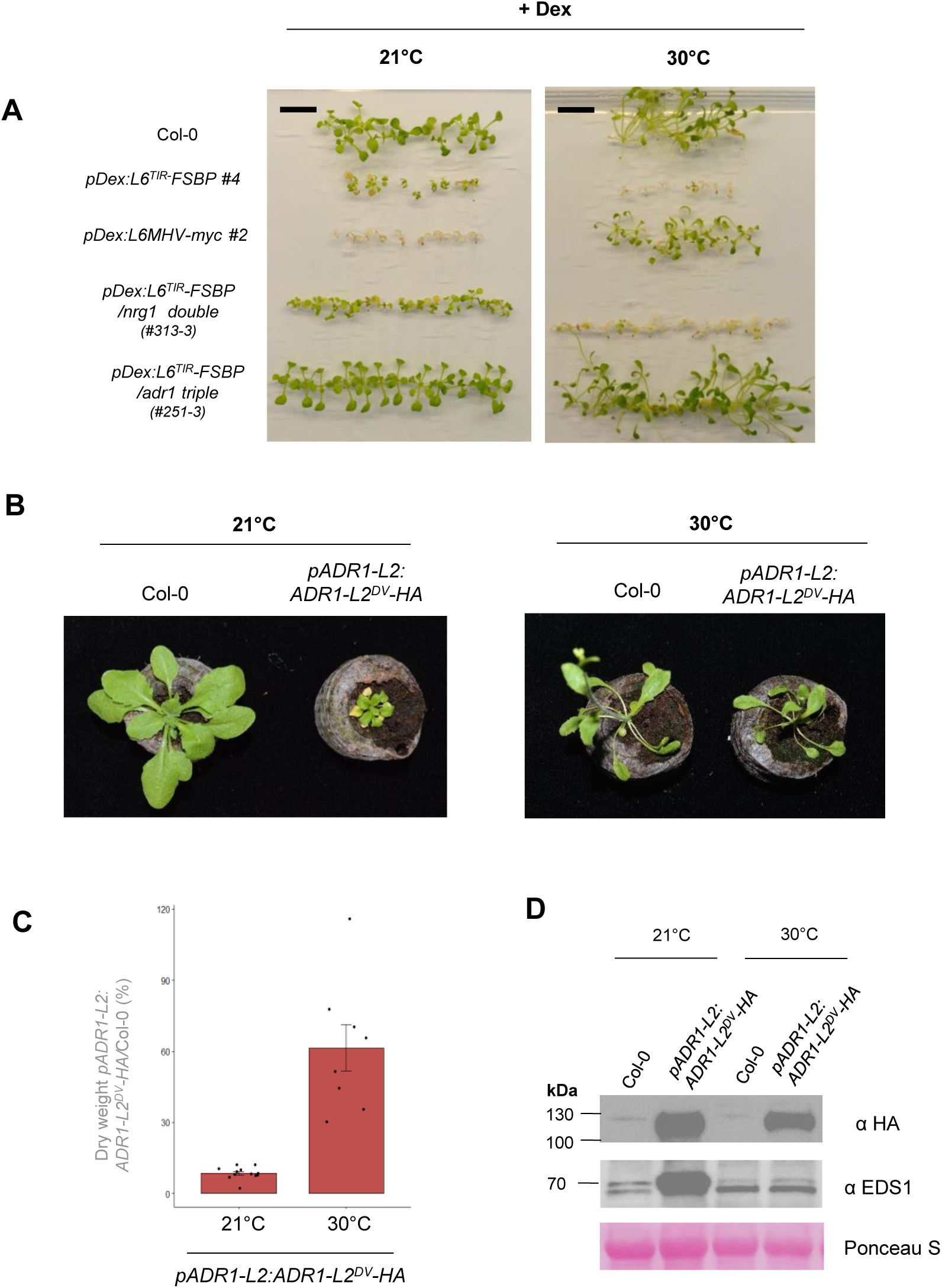
Differential sensitivity of ADR1s-induced signaling to elevated temperature. (A) Cell death phenotype of seedlings carrying *pDex :L6^TIR^* in wild-type, *nrg1 double* or *adr1 triple* mutants, seven days after transfer on a 10 µM Dex medium at 21°C or 30°C. Untransformed wild-type Col-0 was used as a negative control. *pDex :L6MHV-myc line#2* was used as a thermosensitive control. Photos were taken seven days after Dex induction. Black scale bars represent 1 cm. (B) Representative phenotype of three week-old Arabidopsis plant carrying *pADR1-L2:ADR1-L2^DV^-HA* compared to wild-type Col-0 at 21° or 30°C. After germination on MS media, seven day-old seedlings were acclimated for approximately three days on soil pots at 21°C, then transferred and grown for two weeks in climatic chambers set at 21°C or 30°C. (C) Percentage of dry weight ratio of *pADR1-L2:ADR1-L2^DV^-HA* compared to Col-0 plants at 21°C or 30°C as in panel A. Bar plots represent the percentage of mean dry weight ratios of plants carrying *pADR1-L2:ADR1-L2^DV^-HA* / wild-type Col-0 +/- SEM. Individual percentage ratios are represented by black dots (n>8). This experiment was performed twice with similar results. (D) Immunoblot analysis of ADR1-L2^DV^-HA and EDS1 in three week-old transgenics carrying *pADR1-L2:ADR1-L2^DV^-HA* or wild-type Col-0, grown at 21°C or 30°C, using anti-HA and anti-EDS1 antibodies detection, respectively. Total protein load is indicated by Ponceau red staining.

In order to test whether ADR1 family members could individually induce thermotolerant immune signaling, we tested whether the autoimmune phenotype of the previously characterized transgenic line carrying *pADR1-L2:ADR1-L2^DV^-HA* (*60*) was affected by an elevation of temperature. When grown at 21°C, *pADR1-L2:ADR1-L2^DV^-HA* carrying plants exhibit constitutive autoimmune phenotype (dwarf, bushy appearance) with significantly lower dry weight compared to wild-type Col-0, as previously described (*60*)(Figure 4B,C). Surprisingly, we observed that this phenotype was reverted when this mutant was grown at 30°C, despite similar levels of protein accumulation of ADR1-L2^DV^-HA at 21°C or 30°C (Figure 4B-D). Interestingly, EDS1 protein accumulation was significantly enhanced in this *pADR1-L2:ADR1-L2^DV^* line compared to Col-0 at 21°C (Figure 4D), demonstrating that ADR1-L2^DV^-mediated autoimmunity triggers the EDS1 pathway at permissive temperature, like TNL and isolated TIR-induced signaling (Figure 1C). In contrast, EDS1 enhanced accumulation was abolished at 30°C, which supports that ADR1-L2^DV^ -mediated autoimmune signaling is inhibited at 30°C and hence is thermosensitive.

Together, these results indicate that signaling mediated by ADR1s is thermoresilient when induced by isolated *L6^TIR^* but not by autoimmune ADR1-L2^DV^, suggesting a possible differential temperature sensitivity of ADR1 family members.

### Flg22 treatment enhances pDex:TNLs and pDex:isolated TIRs-mediated immune signaling and protein accumulation

Accumulating evidence show that PTI and ETI are tightly interconnected and potentiate each other (*7*). One piece of evidence was that inducible expression of the bacterial effector AvrRps4 did not trigger any cell death in transgenic Arabidopsis line harboring cognate TNL pair RRS1/RPS4, in absence of pathogens. Instead, RRS1/RPS4-mediated specific HR was restored upon PTI eliciting and AvrRPS4 inducing co-treatment (*61*). Our data show that induction of autoimmune TNLs or isolated TIRs expression is sufficient to activate immune responses and cell death in a PAMP-independent manner (Figure 1). This prompted us to test whether this response could be potentiated by PTI or if the signaling pathways downstream of effector recognition does not rely on PTI. For this, we monitored cell death symptoms induced in leaves of adult Arabidopsis lines carrying *pDex:L6^TIR^* or *pDex:L6MHV*, in presence or absence of the PTI elicitor flg22 (Figure 5A). Upon treatment of *pDex:L6^TIR^*and *pDex:L6MHV* lines with mock (DMSO) or flg22 only, infiltrated leaves showed negligible or no symptoms at 2, 3 and 6 days after infiltration, probably due to wounding during infiltration. Dex treatment induced very mild chlorosis two days after infiltration in both *pDex:L6^TIR^* and *pDex:L6MHV* lines. In contrast, most leaves co-treated with Dex and flg22 (Dex+flg22) developed clear chlorotic symptoms from two days after treatment and the difference between Dex only or Dex+flg22 treatment became even clearer at 3 days post treatment, suggesting that cell death induced by full-length L6MHV and isolated L6^TIR^ is accelerated and could be potentiated by PTI (Figure 5A, Figure S10). At 6 days post-treatment, leaves treated with Dex or Dex+flg22 showed, in average, similar level of cell death in all lines. Similar results were observed on at least one independent inducible line carrying *pDex:L6MHV-FSBP (#18)* (Figure S10). Immunoblot analyses showed incremental protein accumulation of both L6^TIR^ and L6MHV at 24h and 48h upon Dex and Dex+flg22 treatments (Figure 5B), as observed in Figure 1C. More surprisingly, we repeatedly observed higher protein accumulation of both L6^TIR^ and L6MHV proteins upon Dex+flg22 co-treatment, compared to Dex-only treatment. However, EDS1 increased accumulation mediated by L6^TIR^ and L6MHV Dex induction remained steady between Dex or Dex+flg22 treatments, suggesting that Dex+flg22 co-treatment potentiate L6^TIR^ and L6MHV protein production but not EDS1 increased accumulation. We obtained similar results with *pDex:RPS4^TIR^*line #F and two independent lines carrying *pDex:RPS4-FSBP (*lines #11 and 14) (Figure S11). Dex+flg22 co-treatment also induced earlier chlorosis/cell death symptoms compared to Dex alone treatment. RPS4^TIR^ and RPS4 full-length protein accumulation was also enhanced in Dex+flg22 compared to Dex alone treatment, while EDS1 accumulation kept steady (Figure S11).

**Figure 5.**
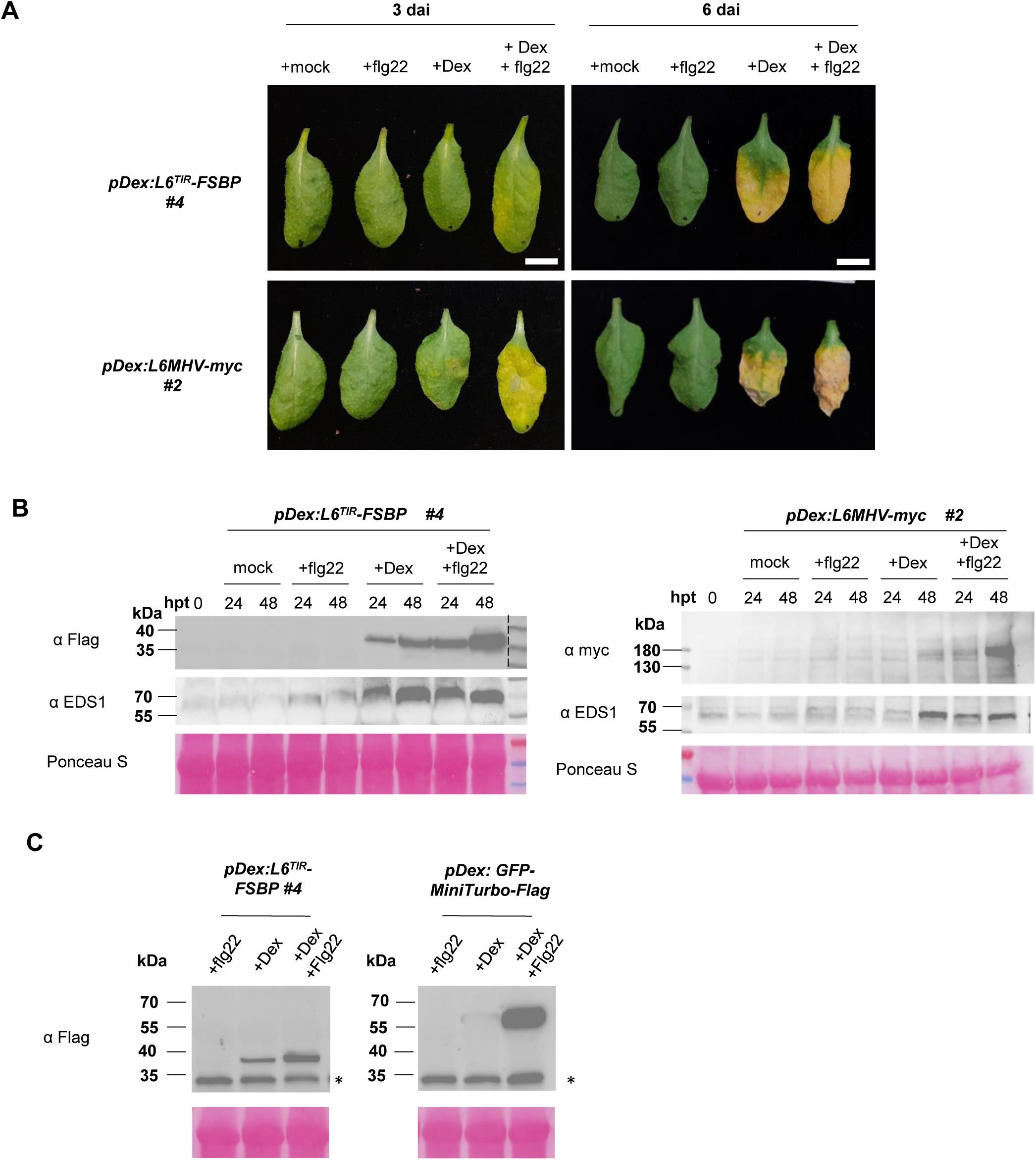
Effect of flg22 treatment on L6TIR- and L6MHV-mediated signaling. (A) Representative cell death symptoms observed in leaves of four week-old Arabidopsis transgenics carrying *pDex:L6^TIR^-FSBP* or *pDex:L6MHV-myc*, infiltrated with a solution of DMSO (“mock”), 100 nM flg22 (“+flg22”), 20 µM Dex (“+Dex”) or a co-treatment of both Dex 20 µM and flg22 100 nM (“+Dex + flg22”). Pictures were taken 3 and 6 days after treatment. (B) Immmunoblot analysis of L6^TIR^-FSBP, L6MHV-myc and EDS1 in leaves of four week-old Arabidopsis transgenics carrying *pDex:L6^TIR^-FSBP* or *pDex:L6MHV-myc*, 24h and 48h post treatment (hpt), using anti-flag, anti-myc or anti-EDS1 antibodies detection, respectively. Three discs from three different plants were collected for L6^TIR^-FSBP detection and five discs from five plants were collected for L6MHV-myc detection. (C) Immmunoblot analysis of L6^TIR^-FSBP and GFP-MiniTurbo-Flag in leaves of four week-old Arabidopsis carrying *pDex:L6^TIR^-FSBP* or *pDex:GFP-Mini-Flag* 24h post treatment, using anti-flag antibodies detection. Background noise is indicated by an asterisk. (B-C) Total protein amount was verified with Ponceau red staining. Immunoblot analyses were performed at least twice with similar results.

Based on these results, we could not conclude whether flg22 treatment potentiates TIR accumulation and hence TIR signaling, or if it has a general impact on the activation of our Dex-inducible system, thus enhancing transcription and protein accumulation and leading to a dose-effect enhanced cell death phenotype. To test this, we used as a control an inducible Arabidopsis line carrying a gene construct that is not involved in immunity (*pDex:GFP-MiniTurbo-flag*). Turbo is a biotin ligase that is used for proximity labelling approaches (*62*). 24h after treatment, plants carrying *pDex:L6^TIR^* or *pDex:GFP-MiniTurbo* accumulated increased level of L6^TIR^ and GFP-MiniTurbo protein, respectively, in leaves treated with Dex+flg22 compared to Dex only treatment (Figure 5C). This demonstrates that flg22 boosts gene expression in our Dex-inducible vector but not native gene expression/protein accumulation such as EDS1 (Figure 5B), and raises caution about data interpretation on PTI/ETI potentiation when using Dex+flg22 co-treatment. As TIR signaling seems to be dose dependent, it is likely that the enhanced cell death phenotype observed upon Dex+flg22 treatment compared to Dex only is due to increased protein production by flg22 instead of a potentiation of TIR signaling by PTI.

## Discussion

NLR activity can be dampened under elevated temperatures, which is concerning in the current climatic context. This observation was previously reported for several TNLs, including Arabidopsis SNC1 or RPS4/RRS1 used in this study (*11, 12, 14*), but the underlying molecular mechanisms remain poorly understood. We showed that immune responses induced in Arabidopsis by different TIR domains isolated from RPS4, SNC1 and flax L6, are maintained at elevated temperatures, unlike those induced by full-length TNLs, suggesting that a raise of temperature impacts TNL receptor functions but not downstream immune signaling.

As signaling units, TIR domains can autonomously induce cell death when overexpressed in absence of pathogens. At 30°C, a non-permissive temperature for most NLRs, we observed that Arabidopsis seedlings expressing RPS4^TIR^, L6^TIR^ and SNC1^TIR^ displayed clear cell death symptoms, which were similar or even stronger than at permissive temperature (21°C), in contrast to autoimmune transgenic lines expressing full-length RPS4 or autoimmune variant L6MHV. To validate this finding, we showed that thermotolerant signaling induced by isolated TIRs share the same signaling components as thermosensitive full-length TNLs. Both isolated TIRs and full-length TNLs promote the accumulation of EDS1 protein and gene expression of major regulators of the SA pathway, such as *CBP60g* and *ICS1,* and require RNLs to trigger signaling. However, EDS1 accumulation and upregulation of *CBP60g* and *ICS1* gene expression is maintained at 30°C in seedlings expressing isolated TIRs but not full-length NLRs, supporting that immune responses induced downstream of TNLs, including the SA pathway, are not affected by an elevation of temperature. This result contrasts with earlier studies reporting that the expression of *CBP60g*, *ICS1* and other SA-related genes was strongly reduced in immune-triggered plants above 28°C (*9, 46*). In the study of Kim et al., the authors showed that elevated temperatures inhibit the SA pathway at its onset, by reducing the number of GUANYLATE BINDING PROTEIN-LIKE 3 (GBPL3) defence-activated biomolecular condensates (GDACs). Consequently, this impairs the recruitment of GBPL3 and associated transcriptional coactivators to the promotor region of *CBP60g* and *SARD1,* which dramatically reduce their expression at 28°C. CBP60g and SARD1 are master transcription factors that regulate the expression of genes involved in SA biosynthesis, such as *ICS1,* which expression is hence turned off under elevated temperature growth conditions (*46*). Interestingly, overexpressing CBP60g or the EDS1/PAD4 complex is sufficient to recover SA signaling and to maintain resistance to virulent and avirulent bacteria at elevated temperature (*45, 46*). Hence, overexpression of immune regulators involved in the SA pathway can bypass the negative impact of temperature on GDAC formation and on plant immunity. Given that the expression of *CBP60g* and *ICS1* is dramatically reduced at 30°C upon activation of thermosensitive NLRs, but not when isolated TIR domains are overexpressed, our results demonstrate that activating downstream TIR signaling is also sufficient to bypass the negative impact of temperature on the SA sector. It also supports that the activation status of full-length receptors is negatively impacted by warmer temperature (above 28°C) whereas downstream signaling, including the SA pathway, is thermostable. Hence, it is possible that reduced expression of *CBP60g* and *ICS1* may be controlled by NLR activation status rather than by temperature directly. Alternatively, TNL signaling may activate the expression of *CBP60g* in a GDAC-independent manner. NLRs, and in particular TNLs, widely contribute to different levels of plant immunity. While NLRs were first identified as effector receptors to trigger ETI, they also contribute to amplify PRR-mediated immunity (*7, 63, 64*). For example, global disruption of NLR homeostasis dramatically reduce PTI (*63*). Hence, if NLRs function upstream of GDAC formation, temperature vulnerability of the SA pathway may be explained by a collective inactivation of thermosensitive NLRs under elevated temperature. It would be interesting to monitor the formation of GDACs as well as the recruitment of GBPL3 and coactivators to the *CBP60g* promoter upon expression of isolated TIR domains or full-length NLRs to decipher whether TNL-induced expression of *CBP60g* is GDAC-dependent or -independent, especially under elevated temperature.

All TNLs characterized so far require RNLs to induce immune signaling. RNLs function redundantly and contribute differentially to cell death or basal defense and gene reprogramming (*26, 42, 65*). By using Arabidopsis lines expressing isolated TIR domains in different RNLs subgroup mutant backgrounds, we showed that immune signaling induced by isolated TIR domains depends on RNLs, just as effector-dependent or –independent full-length TNL-induced signaling. Here, we showed by monitoring macroscopic cell death, that L6^TIR^ and SNC1^TIR^ signaling mainly depend on the ADR1 family to translate cell death, which is consistent with previously reported requirement of ADR1s for *snc1* autoimmune phenotype (*42*). For RPS4^TIR^-mediated immune responses, each RNL subgroup seems to compensate for the absence of the other as RPS4^TIR^ -mediated autoimmunity (severe stunting to cell death) was still visible in both *nrg1 double* and *adr1 triple* mutants, whereas it was fully abolished in the quintuple *helperless* mutant. These results slightly differ from previous studies, in which RPS4/RRS1-mediated cell death requires NRG1s whereas the ADR1 family supports disease resistance and pathogen multiplication restriction (*26*). Importantly, our results show that thermotolerant flax L6^TIR^ share the same genetic requirements as the corresponding thermosensitive full-length receptor L6MHV, further supporting that this shared pathway is maintained under elevated temperature only when induced by isolated TIR and not by full-length TNL. These data also suggest ADR1s-mediated signaling is thermostable when activated by isolated TIRs. Indeed, we showed that L6^TIR^ can still induce immune signaling in *nrg1 double* mutant at 30°C but not in *adr1 triple* mutant, supporting our primary hypothesis that ADR1s can translate TIR signaling at elevated temperature. However, the autoimmune phenotype mediated by one of the ADR1 family member in the *pADR1:ADR1-L2^DV^-HA* line at 21°C was reverted at 30°C, suggesting that ADR1-L2 *^DV^* is thermosensitive. Interestingly, EDS1 enhanced protein accumulation, which correlates with this autoimmune phenotype, is also dramatically reduced at 30°C in this line. This is consistent with previous studies showing that *pADR1:ADR1-L2^DV^-HA-*mediated autoimmunity genetically requires *EDS1*, possibly to positively regulate SA accumulation via a feedback loop (*60, 66*), which seems to be inhibited at 30°C (this study). The underlying mechanisms will need to be further investigated to fully understand the differential temperature sensitivity of ADR1s. Another intriguing question is whether ADR1 family members have differential sensitivity to elevated temperature or whether *ADR1-L2^DV^-HA* autoimmune variant activates a separate pathway that is thermosensitive. ADR1-L2^DV^-mediated autoimmunity signals through TNL *SADR1* (Suppressor of *ADR1-L2 1*), which is dispensable for signaling induced by other TNLs (*66*). It remains to be determined whether, as for many full-length TNLs, SADR1 function is inhibited at 30°C too. Alternatively, it is possible that activation of ADR1s in the pDex: L6^TIR^ /*nrg1 double* line may induce a SADR1-independent pathway, which is resilient to elevated temperature, explaining why we could still observe cell death at 30°C.

Overall, our results point towards the thermoresilence of immune signaling induced by isolated TIR domains, in contrast to corresponding full-length TNLs, which may be explained by a simpler architecture of isolated TIRs. Naturally-occurring truncated NLRs, such as, lack some of the canonical NLR domains (LRR and/or NB domain) but contain a TIR domain. TIR-only or TIR-NB coding genes are widely represented in Arabidopsis genome and massive transcriptional induction of these genes upon immune stimuli indicate they are broadly involved in plant immunity (*55, 67–69*). Interestingly, we found that two previously described TIR-containing truncated TNLs (RBA1 and TN2) induce thermostable immune signaling. Hence, it will be of great interest to further characterize the function of such non canonical TIR-containing proteins and assess whether they can induce thermotolerant disease resistance. Given that NLRs are broadly used in crop breeding programs for disease resistance, this work underlines the need to further investigate how the function of TNLs or more generally NLRs is impacted by temperature stress. Canonical sensor TNLs are modular proteins which activation is finely regulated via intra- and inter-domain interactions as well as interactions with chaperones proteins (*18, 70, 71*). Therefore, it is likely that such complex mode of regulation may be affected by environmental changes such as temperature rise, which in turn may turn off TNL-induced enzymatic activities and downstream signaling.

Several studies have suggested that NLRs could function at the interface between biotic and abiotic stress responses (*72–74*), pointing to a role as mediators of growth-defense trade-offs. Hence, it will be important in the future to further investigate how NLRs function is modulated by different abiotic stresses.

### Limitations of the study

- By using inducible transgenic lines expressing autoimmune TNLs or isolated TIRs, this study allowed us to bypass effector recognition to focus on TIR-activated downstream signaling. However, this system revealed some limitations as the signaling output monitored (cell death) is dose-dependent. To deepen our understanding on the impact of temperature stress and the differential requirement of RNL family members for TIR signaling, further RNA-seq analyses would be necessary to obtain a full picture of the plant response in these conditions.
- Our Dex-inducible vector is sensitized by PTI elicitor flg22. This allowed us to raise attention to the interpretations PTI/ETI interconnections studies when using such system. Hence, it prevented us to conclude on whether PTI may potentiate effector-independent TIR signaling.
- Our data were obtained upon a single temperature stress in controlled conditions. It will be important in the future to further explore the plant immune response upon combined biotic and abiotic stresses (temperature, drought, CO2 concentration…), to better mimic predicted climate change conditions.

## METHODS DETAILS

### Plasmid constructions

All plasmid constructs used for Arabidopsis stable transformation were generated by Gateway cloning (GWY, Invitrogen). PCR products flanked by attB sites were recombined into pDONR207 (Invitrogen) and then into dexamethasone inducible pOpOff2-derived destination vectors pDex:GWY-FSBP, pDex:GWY-YFPv or pDex:GWY-MiniTurbo-3Flag (*53, 75*). Generation of pDex:GWY-FSBP was previously described in (*53*). The FSBP tag consist of a triple flag epitope (F) fused with a streptavidin-binding peptide (SBP) (*53*). pDex:GWY-YFPv or pDex:GWY-MiniTurbo-3Flag were modified from pOpOff2(Kan) GWY vector, which was kindly provided by Chris Helliwell (CSIRO, Canberra). This vector was lacking the hairpin GWY cassette (as described in (*76*)) and instead contained a simple GWY cassette but no terminator. Hence, a PCR fragment containing either the DNA sequence of the yellow fluorescent protein venus (YFPv)(*77*) or the MiniTurbo (*62*) with an in frame triple flag epitope sequence, followed by the 35s terminator sequence, flanked with *KpnI/Pme1* restriction sites, were used to build pDex:GWY-YFP or pDex:GWY-MiniTurbo-3Flag by restriction/ligation using pOpOff2(Kan)-GWY as the vector backbone. All plasmids and constructs were verified by sequencing. All plasmids, constructs and primers used in this study are listed in Tables S2 and S3.

### Plant growth conditions

Arabidopsis seeds were sown on MS media supplemented with antibiotics when needed, for transgenic lines selection. After 24h vernalization, MS plates were transferred in growth chambers under 16h photoperiod, 21°C conditions for eight to ten days. For seedling assays, eight to ten day-old seedlings were transferred to regulated climatic chambers (Memmert ®) set at 21°C or 30°C, under 8h photoperiod, after Dexamethasone treatment (see details in Dex-induction section below). For adult plant assays, eight day-old seedlings were transferred on soil pots (Jiffy®) and grown for two to three weeks in growth chambers under 8h photoperiod conditions at 21°C, and transferred to climatic chambers (Memmert ®) set at 21°C or 30°C, under 8h photoperiod, after treatment (dexamethasone or flg22, see details in Dex-induction section below).

### Plant material and transgenic lines screening

Arabidopsis Col-0, and mutants in Col-0 background (*eds1-2* (*78*) or *RNL* mutants (*26*), were transformed by floral-spraying or floral-dipping using *Agrobacterium tumefaciens* strains carrying *pDex:TIRs* or *pDex:TNLs* constructs. Primary transformants were selected on MS media supplemented with hygromycin (50 µg/ml) and were individually PCR genotyped to verify the presence of the transgene or mutations in *RNLs* by using Thermo Scientific™ Phire plant direct PCR kit as previously described (*53*). Used primers for PCR genotyping are listed in Table S3. Western-blot analyses were further performed on F2 progenies to verify protein accumulation of TIR or TNL constructs, 24h after dexamethasone induction of 8 day-old seedlings. *pADR1:ADR1-L2^DV^-HA* line (*60*) was kindly provided by F. El Kasmi. All transgenic lines used in this study are listed in Table S1. For dry weight measurement, plants were individually harvested, dried in paper pockets at 37°C and weighed 3 days after.

### Cell death assays upon dexamethasone induction and/or flg22 treatment

For seedling assays, seven to ten day-old seedlings were transferred on MS media plates supplemented with Dexamethasone (10µM) or DMSO (equivalent volume as Dexamethasone) and transferred to regulated climatic chambers (Memmert ®) set at 21°C or 30°C, under 8h photoperiod.

For adult plants assays, 8 day-old seedlings were transferred on soil pots (Jiffy®) and grown for two to three weeks in a growth chamber under 8h photoperiod conditions at 22°C. Three to five leaves of three to five week-old plants were infiltrated with a needle-less syringe-with solutions containing 20 µM Dexamethasone or mock solution (equivalent volume of DMSO as Dexamethasone), a combination of flg22 peptide (100nM) and DMSO; or a combination of 20 µM Dexamethasone and 100nM flg22 solutions. For temperature assays, plants were transferred to regulated climatic chambers (Memmert®) set at 21°C or 30°C, under 8h photoperiod upon Dex treatment. The apparition of cell death symptoms was monitored every day.

### Immunoblot analyses

For seedling assays, total protein extraction was performed on 10 eight day-old seedlings collected at different time points after Dexamethasone treatment. For adult plant assays, three or five 6mm leaf discs were harvested from three to five different plants per line carrying *pDex:RPS4^TIR^* or *pDex:L6^TIR^*, or lines carrying *pDex:RPS4* or *pDex:L6MHV*, respectively, 24h and 48h after treatment. For ADR1-L2^DV^-HA detection, 7 leaf discs (diameter 1cm) were collected from 3 week-old plants grown at 21°C or 30°C. Plant material expressing TIR domains were ground and directly resuspended in 100µl of loading Laemmli buffer. Plant material expressing TNLs or RNLs were ground and resuspended in 100µl extraction buffer [150mM Tris-HCl pH7.5, 150mM NaCl, 1mM EDTA, NP-40 1%, Plant protease inhibitor cocktail (SIGMA) 1%, 10 mM DTT, 10 µM MG132], centrifuged for 5min at 4°C to remove cell debris and 50µl of the supernatant was combined with 50µl loading Laemmli buffer (0.125 M Tris HCl PH7.5, 4% SDS, 20% Glycerol, 0.2M DTT, 0.02% Bromophenol blue) for western-blot analysis. After denaturation at 95°C for three minutes, samples were then loaded on sodium dodecyl sulfate polyacrylamide gels for electrophoresis and transferred to nitrocellulose membranes which were blocked with 5% skimmed milk. Membranes were stained using Ponceau red staining to verify total protein loading. Membranes were further probed with anti-flag-peroxidase (HRP) (Sigma, SAB4200119) (dilution ratio: 1/5000), anti-Myc-peroxidase (clone 9E10; Roche) (dilution ratio: 1/5000), or rabbit *anti-EDS1* (kindly provided by J. E. Parker) (dilution ratio: 1/500) followed by incubation with a secondary goat anti-rabbit antibodies (Bio-Rad) conjugated to peroxidase (dilution ratio: 1/10000). Protein detection was performed using Bio-Rad Chemidoc™ Imaging system with the Bio-Rad Clarity Western ECL Substrate or Bio-Rad Clarity Max Western ECL Substrate.

### RNA extraction and qPCR analyses

Total RNA extraction was performed on 10 eight day-old seedlings or three 6mm leaf discs from three different four week-old plants at 24h and 48h after treatment. Total RNA was extracted using the Macherey Nagel Nucleospin RNA Plus kit following manufacturer’s instruction. Total RNA was then subjected to DNAse treatment according to the Invitrogen™ Ambion™ TURBO DNA-free Kit instructions. 1µg total RNA was used for reverse transcriptase reactions. RT-qPCR reaction mix was prepared with SYBR green (Takyon™ No ROX SYBR 2X MasterMix blue dTTP) and run on Bio-Rad CFX Opus 384 Real-Time PCR instrument. Used primers are listed in Table S3.

### Quantification and statistical analysis

For RT-qPCR data, mean ΔCt were calculated from three technical replicates from two to four biological replicates (as indicated in figure legends). *At1G13320* and *At5G15710* were used as internal control genes. Log2(2^-^ΔCt) values were used to determine the significance of difference in gene expression between the different tested conditions, and were represented as boxplots using R studio. A two-ways ANOVA was used when the distribution of the data fitted assumptions of normality and homogeneity of variances, and was followed by a post-hoc pairwise comparison Tukey test.

## Acknowledgments

This research was set within the framework of the “Laboratoires d’Excellence (LABEX)” TULIP (ANR-10-LABX-41) and of the “École Universitaire de Recherche (EUR)” TULIP-GS (ANR-18-EURE-0019). HD was supported by a PhD scholarship funded by the French Ministry of National Education and Research. MB was supported by a research grant funded by INRAE (Plant health department) (PIMS). LD was supported by a research grant funded by the Agence Nationale de la Recherche ANR-18-CE20-0015. We thank Dr F. El Kasmi (Tubingen University, Germany) for sending Arabidopsis mutant lines (*RNLs*) and ADR1 plasmid constructs, and for interesting discussions.

## Author contributions

HD and MB designed and performed experiments. CB generated transgenic plant material. HD, LD, and MB analyzed data. HD and MB wrote the manuscript. HD, LD, and MB edited the manuscript.

## Supplementary data

### Supplementary figures titles and legends

Figure S1: Dex-induced seedling cell death phenotype and protein accumulation of independent transgenic lines used in this study.

Figure S2. Effect of temperature on Dex-induced cell death phenotype and protein accumulation in leaves of independent transgenic lines used in this study.

Figure S3. SNC1^TIR^ induces autoimmune symptoms at 21°C and 30°C.

Figure S4. Naturally occurring TIR-containing truncated TNL proteins induce autoimmune symptoms at 30°C.

Figure S5: L6^TIR^-mediated immunity depends on EDS1 in Arabidopsis.

Figure S6. Seedling phenotype and TIR protein accumulation in independent Arabidopsis lines carrying *pDex:RPS4^TIR^-FSBP* in different RNL mutant backgrounds (*nrg1 double, adr1 triple, helperless*).

Figure S7. Seedling phenotype and TIR protein accumulation in independent Arabidopsis lines carrying *pDex:SNC1^TIR^-FSBP* in different RNL mutant backgrounds (*nrg1 double, adr1 triple, helperless*).

Figure S8. Seedling phenotype and TIR protein accumulation in independent Arabidopsis lines carrying *pDex:L6^TIR^-FSBP* in different RNL mutant backgrounds (*nrg1 double, adr1 triple, helperless*).

Figure S9. Seedling phenotype and *L6MHV* transcript levels in independent Arabidopsis lines carrying *pDex:L6MHV-FSBP* in different RNL mutant backgrounds (*nrg1 double, adr1 triple, helperless*).

Figure S10. Effect of flg22 co-treatment on Dex-induced cell death in independent Arabidopsis lines carrying *pDex:L6MHV-myc or pDex:L6MHV-FSBP*.

Figure S11. Effect of flg22 co-treatment on Dex-induced cell death in Arabidopsis lines carrying *pDex:RPS4^TIR^-FSBP* and *pDex:RPS4-FSBP*.

## Supplemental information

Table S1: List of transgenic lines used in this study

Table S2: List of constructs used in this study

Table S3: List of primers used in this study

**References associated to supplementary data**

## RESOURCE AVAILABILITY

### Lead contact

Further information and requests for resources and reagents used in this study should be directed to and will be fulfilled by the lead contact, Maud Bernoux.

