## supplemental data for "Immune signaling induced by plant Toll/interleukin-1 receptor (TIR) domains is thermostable"

FigS1

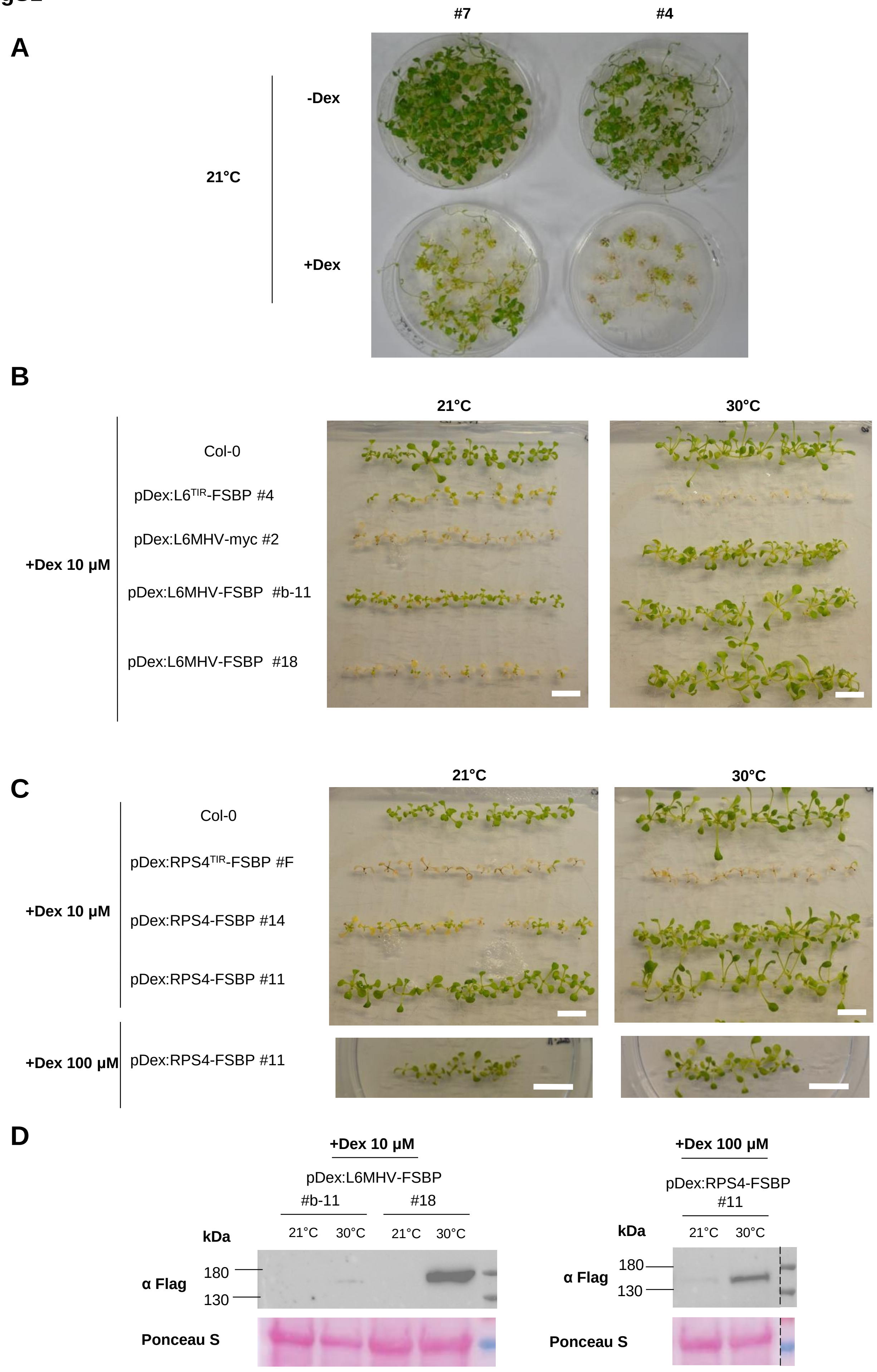
A
#7
#4
21°C
-Dex
+Dex
B
21°C
30°C
+Dex 10 µM
Col-0
pDex:L6TIR-FSBP #4
pDex:L6MHV-myc #2
pDex:L6MHV-FSBP #b-11
pDex:L6MHV-FSBP #18
C
21°C
30°C
+Dex 10 µM
Col-0
pDex:RPS4TIR-FSBP #F
pDex:RPS4-FSBP #14
pDex:RPS4-FSBP #11
+Dex 100 µM
pDex:RPS4-FSBP #11
D
+Dex 10 µM
pDex:L6MHV-FSBP
#b-11
#18
kDa
21°C
30°C
21°C
30°C
180
130
α Flag
Ponceau S
+Dex 100 µM
pDex:RPS4-FSBP
#11
kDa
21°C
30°C
180
130
α Flag
Ponceau S

**Figure S1: Dex-induced seedling cell death phenotype and protein accumulation of independent transgenic lines used in this study.** A. Cell death phenotype of two independent transgenic Arabidopsis lines carrying *pDex:L6*$^{TIR}$*-FSBP*. Seedlings (10 days-old) were transferred to non-inducing (DMSO, -Dex) or 10µM Dex-containing (+Dex) MS media at 21°C. Photos were taken 20 days after transfer. B. Cell death phenotype of two independent Arabidopsis transgenic lines carrying *pDex:RPS4-FSBP*, with wild-type Col-0, and *pDex:RPS4*$^{TIR}$*-FSBP* line #F (Bernoux et al. 2023) as controls. One week-old seedlings were transferred to 10µM and/or 100µM Dex-containing media and moved to climatic chambers at 21°C or 30°C for seven days. C. Cell death phenotype of two independent Arabidopsis transgenic lines carrying *pDex:L6MHV-FSBP*, with wild-type Col-0, *pDex:L6*$^{TIR}$*-FSBP* line *#4* and *pDex:L6MHV-3myc* line *#2* (Bernoux et al. 2023) as controls. One week-old seedlings were transferred to 10µM Dex-containing media and moved to climatic chambers at 21°C or 30°C for seven days. D. Immunoblot analysis of RPS4-FSBP and L6MHV-FSBP in indicated independent transgenic lines 24h after Dex induction at 21°C and 30°C, using anti-Flag antibodies. Total protein load is indicated by Ponceau red staining.

**FigS2**

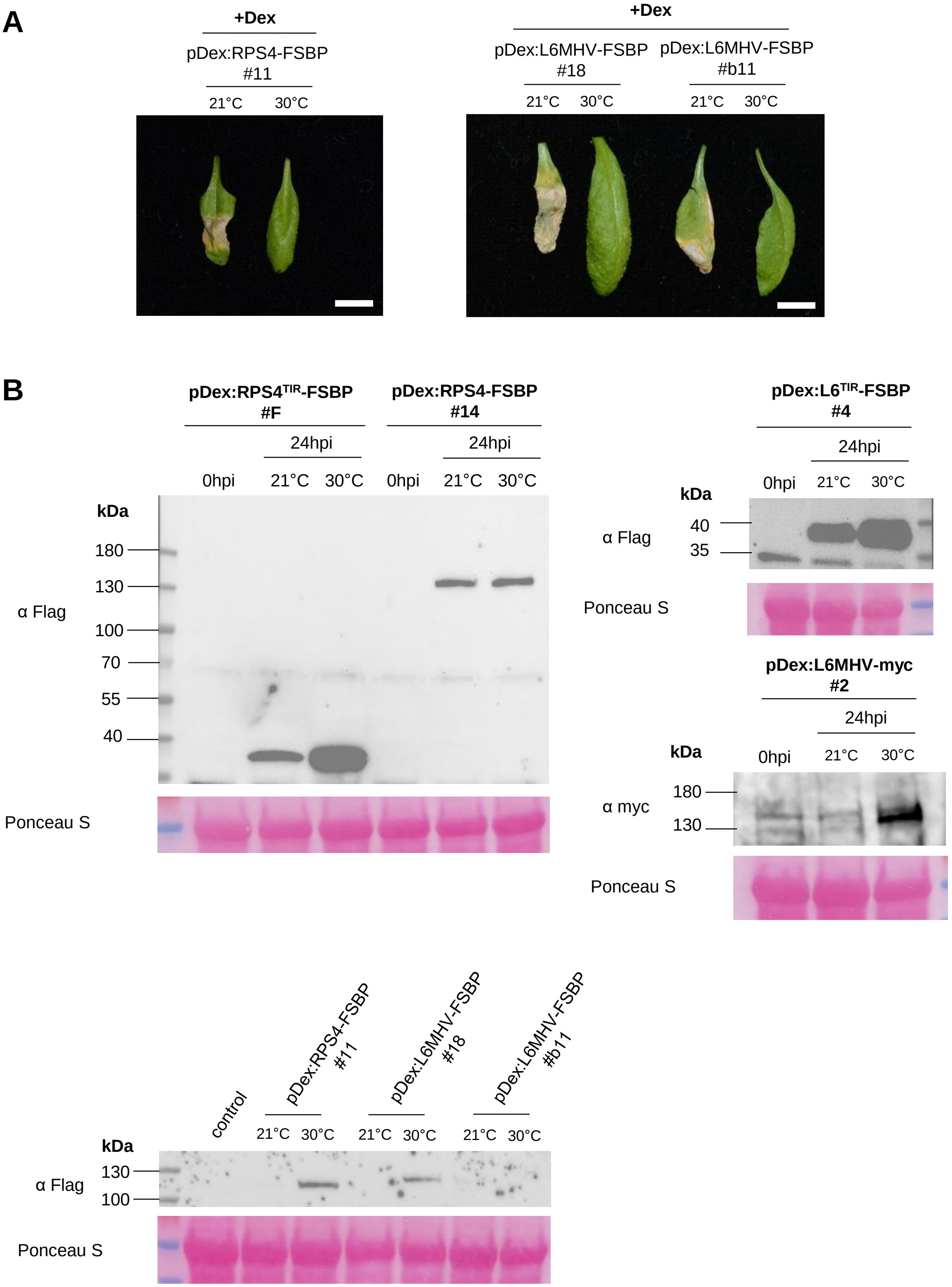


**Figure S2. Effect of temperature on Dex-induced cell death phenotype and protein accumulation in leaves of independent transgenic lines used in this study.** A. Cell death phenotype of independent Arabidopsis transgenic lines carrying *pDex:RPS4-FSBP* or *pDex:L6MHV-FSBP*, seven days after infiltration of 3 weeks-old plant leaves with a 20 µM Dex solution. Plants were moved to climatic chambers at 21°C or 30°C after leaf infiltration. White scale bars indicate 1 cm. B. Immunoblot analysis of L6$^{TIR}$-FSBP, L6MHV-myc, RPS4$^{TIR}$-FSBP and RPS4-FSBP 24h after leaf infiltration of independent transgenic lines with a 20 µM Dex solution at 21°C or 30°C.Total protein load is indicated by red Ponceau staining.

**FigS3**

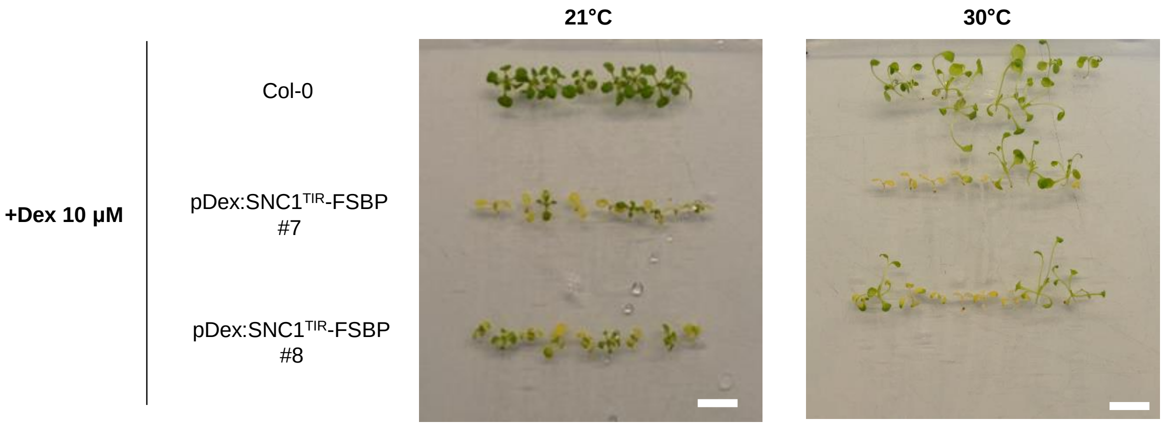


**Figure S3. SNC1$^{TIR}$ induces autoimmune symptoms at 21°C and 30°C.** Cell death phenotype of transgenic seedlings from two independent lines carrying *pDex:SNC1$^{TIR}$-FSBP*. Eight day-old seedlings were transferred to 10µM Dex-containing (+Dex) MS media and moved to climatic chambers at 21°C or 30°C. Wild-type Col-0 seedlings were used as control. Photos were taken 7 days after Dex induction.

**FigS4**

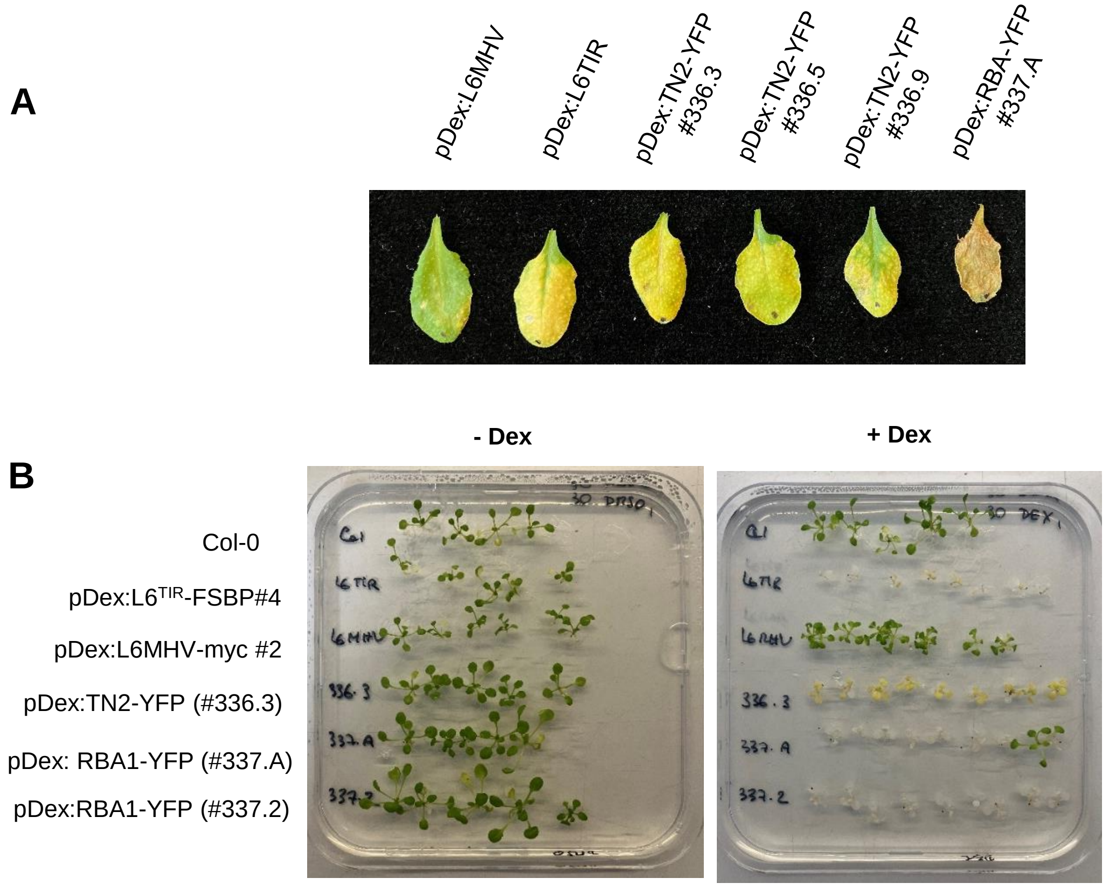


**Figure S4. Naturally occurring TIR-containing truncated TNL proteins induce autoimmune symptoms at 30°C.** (A-B) Cell death phenotype of independent Arabidopsis transgenic lines carrying *pDex:TN2-YFP* or *pDex:RBA1-YFP* after Dex induction. Wild-type Col-0, thermotolerant *pDex:L6$^{TIR}$-FSBP* line *#4* and thermosensitive *pDex:L6MHV-3myc* line *#2* were used as controls. A. Leaves of four week-old T1 individuals were infiltrated with a 20 µM Dex solution and moved to climatic chambers at 30°C. Photos were taken 3 days after Dex induction. B. Eight day-old seedlings (T2) were transferred to non-inducing (DMSO, -Dex) or 10µM Dex-containing (+Dex) MS media and moved to climatic chambers at 30°C. Photos were taken 7 days after Dex induction.

#### FigS5

Col-0

pDex:$L6^{TIR}$-FSBP
(line #4)

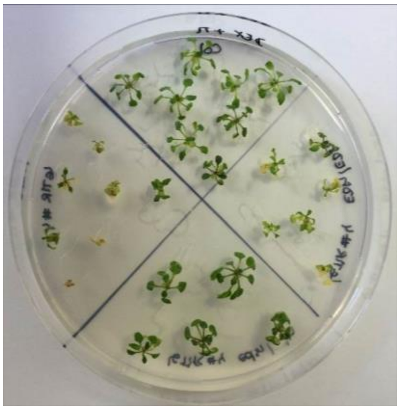

pDex:L6TIR-FSBP
X *eds1-2*
(*EDS1/EDS1* or *EDS1*/*eds1)*

pDex:$L6^{TIR}$-FSBP
X *eds1-2*
*(eds1/eds1)*

**Figure S5: $L6^{TIR}$-mediated immunity depends on EDS1 in Arabidopsis.**

Cell death phenotype of transgenic Arabidopsis seedlings carrying *pDex:$L6^{TIR}$ –FSBP* in *eds1-2* mutant background. F2 individuals from pDex:$L6^{TIR}$ –FSBP (line #4) x *eds1-2* cross were PCR screened to select seedlings homozygous for *EDS1* deletion (*eds1/eds1*) or displaying heterozygous or homozygous wild-type *EDS1 (EDS1/eds1 or EDS1/EDS1)* as described in Bernoux et al. (2023). Ten days-old seedlings were transferred to 10µM Dex-containing media. Photos were taken 12 days after Dex induction.

**FigS6**

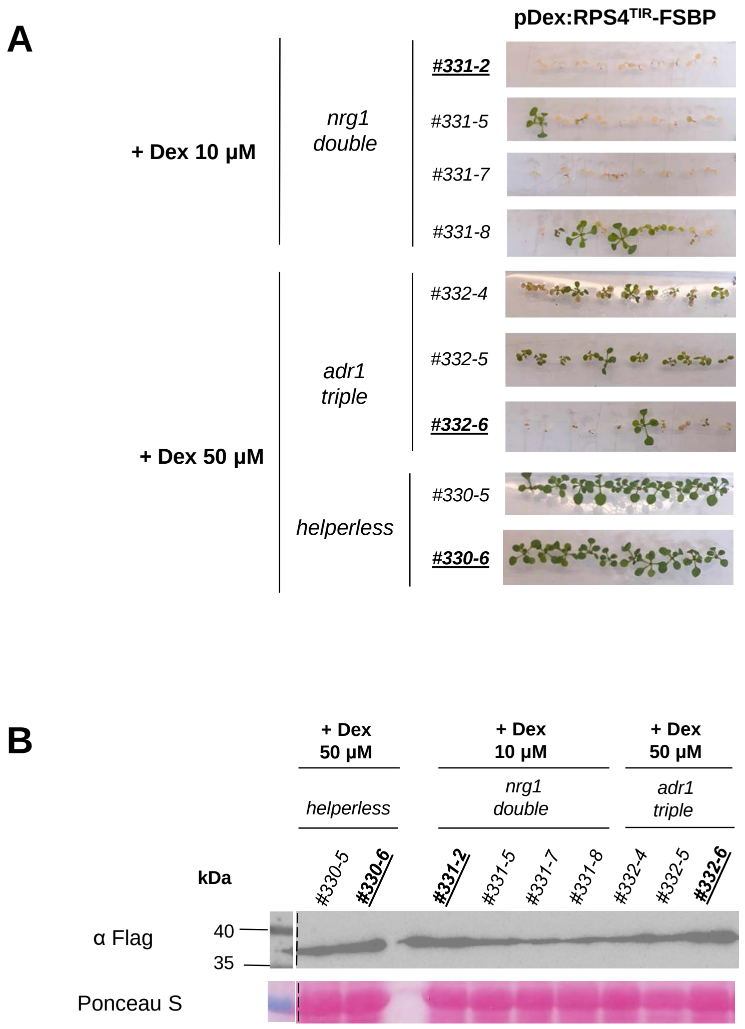


**Figure S6. Seedling phenotype and TIR protein accumulation in independent Arabidopsis lines carrying *pDex:RPS4$^{TIR}$-FSBP* in different RNL mutant backgrounds (*nrg1 double, adr1 triple, helperless*).** A. Cell death phenotype of two weeks-old seedlings from independent transgenic lines carrying *pDex:RPS4TIR-FSBP* seven days after transfer on 10 or 50 µM Dex-containing MS media. B. Immunoblot analysis of RPS4$^{TIR}$-FSBP in indicated independent transgenic lines 24h after Dex induction at 21°C, using anti-Flag antibodies. Total protein load is indicated by Ponceau red staining. For each panel, bold and underlined text represents the lines used for Figure 3.

**FigS7**

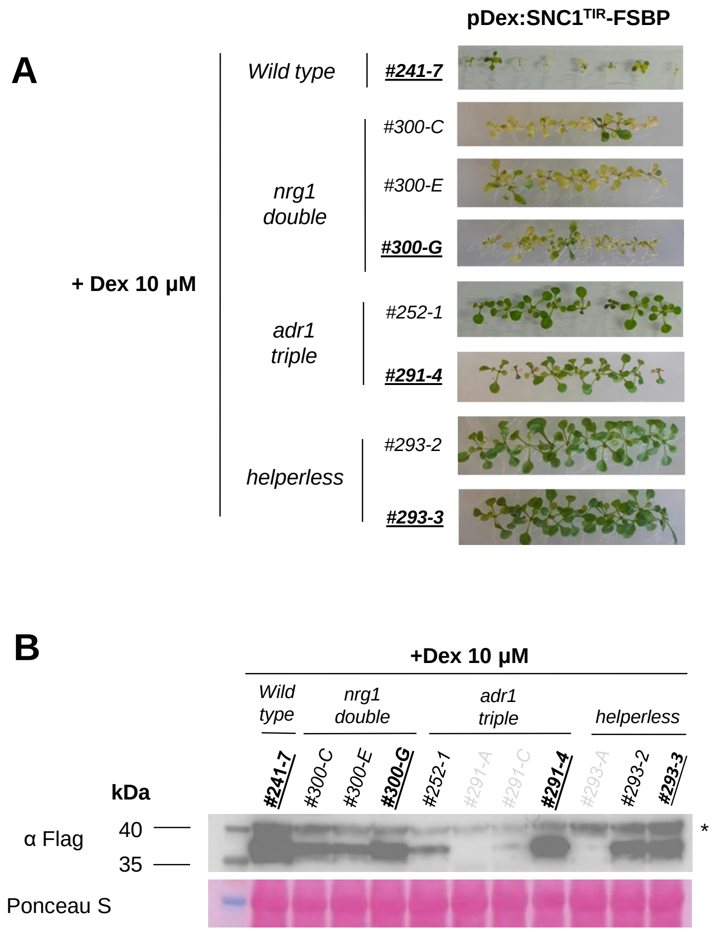


**Figure S7**. **Seedling phenotype and TIR protein accumulation in independent Arabidopsis lines carrying *pDex:SNC1^TIR^-FSBP* in different RNL mutant backgrounds (*nrg1 double, adr1 triple, helperless*).** A. Cell death phenotype of two weeks-old seedlings from independent transgenic lines carrying *pDex:SNC1$^{TIR}$-FSBP* seven days after transfer on 10µM Dex-containing MS media. B. Immunoblot analysis of SNC1$^{TIR}$-FSBP in indicated independent transgenic lines 24h after Dex induction at 21°C, using anti-Flag antibodies. Total protein load is indicated by Ponceau red staining. For each panel, bold and underlined text represents the lines used for Figure 3. Lines number in light grey colour in panel B display no protein accumulation and are therefore not shown in panel A. Non specific bands are indicated by an asterisk.

**FigS8**

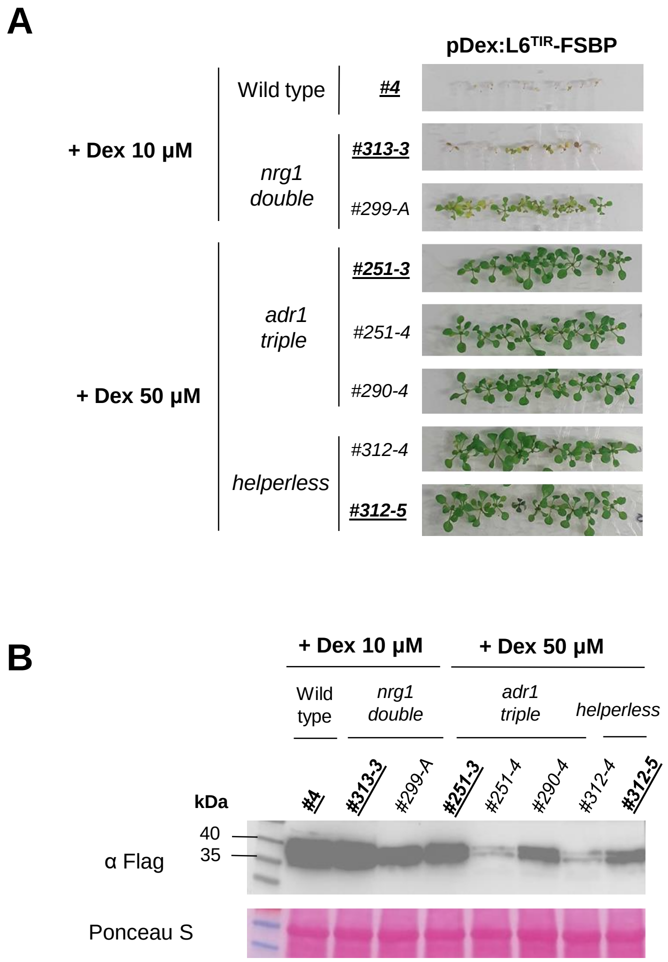


**Figure S8. Seedling phenotype and TIR protein accumulation in independent Arabidopsis lines carrying *pDex:L6$^{TIR}$-FSBP* in different RNL mutant backgrounds (*nrg1 double, adr1 triple, helperless*)**. A. Cell death phenotype of two weeks-old seedlings from independent transgenic lines carrying *pDex:L6$^{TIR}$-FSBP* seven days after transfer on 10 or 50µM Dex-containing MS media. B. Immunoblot analysis of L6$^{TIR}$-FSBP in indicated independent transgenic lines 24h after Dex induction at 21°C, using anti-Flag antibodies. Total protein load is indicated by Ponceau red staining. For each panel, bold and underlined text represents the lines used for Figure 3.

**FigS9**

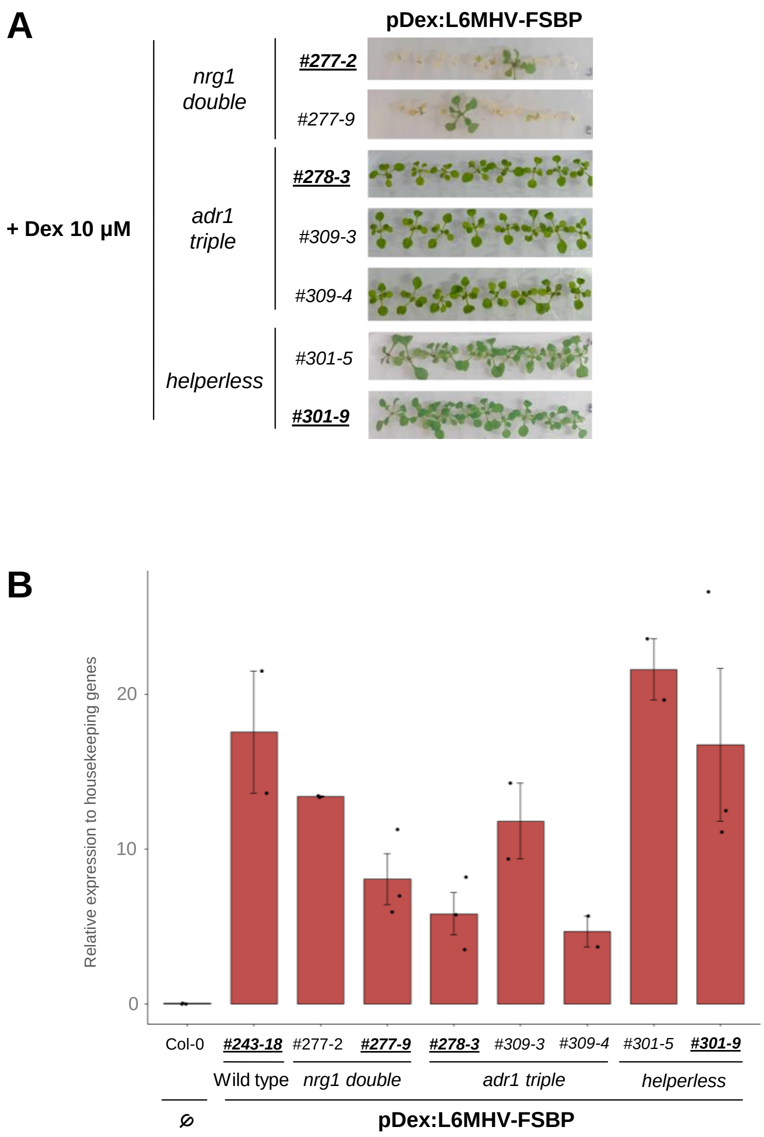


**Figure S9. Seedling phenotype and *L6MHV* transcript levels in independent Arabidopsis lines carrying *pDex:L6MHV-FSBP* in different RNL mutant backgrounds (*nrg1 double, adr1 triple, helperless*).** A. Cell death phenotype of two weeks-old seedlings from independent transgenic lines carrying *pDex:L6MHV-FSBP* seven days after transfer on 10µM Dex-containing MS media. B. RT-qPCR analysis of *pDex:L6MHV-FSBP* transgene in indicated independent transgenic lines 24hours after Dex induction at 21°C. Untransformed wild-type Col-0 was used as a negative control. *pDex:L6MHV-FSBP* expression was normalized relatively to the expression of two housekeeping genes (*At1G13320* and *At5G15710*). Bar plots represent means +/-SEM obtained from two or three independent biological replicates which are represented by black dots. Bold and underlined text represents the lines used for Figure 3.

FigS10

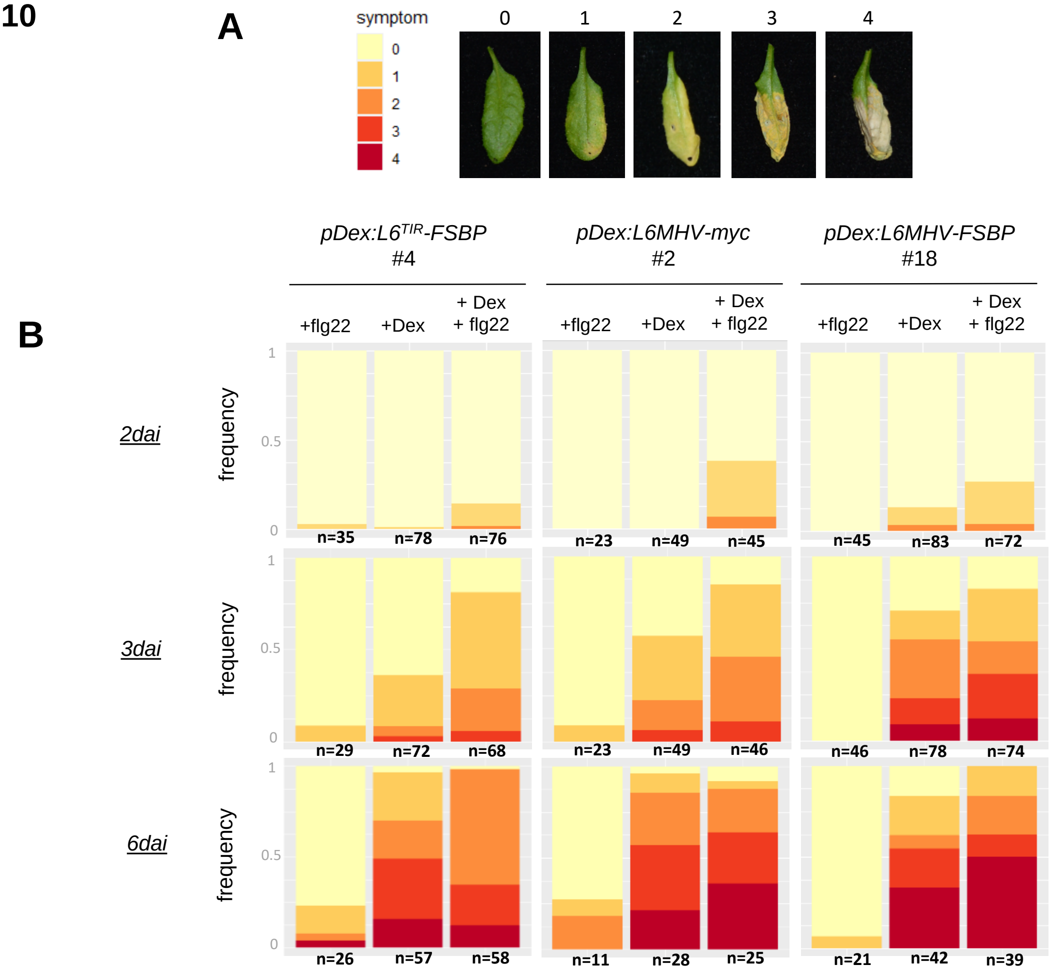


**Figure S10. Effect of flg22 co-treatment on Dex-induced cell death in independent Arabidopsis lines carrying *pDex:L6MHV-myc or pDex:L6MHV-FSBP*. A.** Cell death scoring scale (0 : no symptom, to 4 : complete necrosis. B. Representation of cell death scores as a percentage of each score (color-coded as indicated in [A]), in n>20 leaves of three week-old independent Arabidopsis lines carrying *pDex:L6*$^{TIR}$ or *pDex:L6MHV* (2, 3 or 6 days after infiltration with 100 nM flg22, 20µM Dex, or Dex + flg22 solutions). Numbers of scored leaves per treatment is indicated at the bottom of the graphs for each condition. Data presented in (B) are the result of at least three independent experiments.

### FigS11

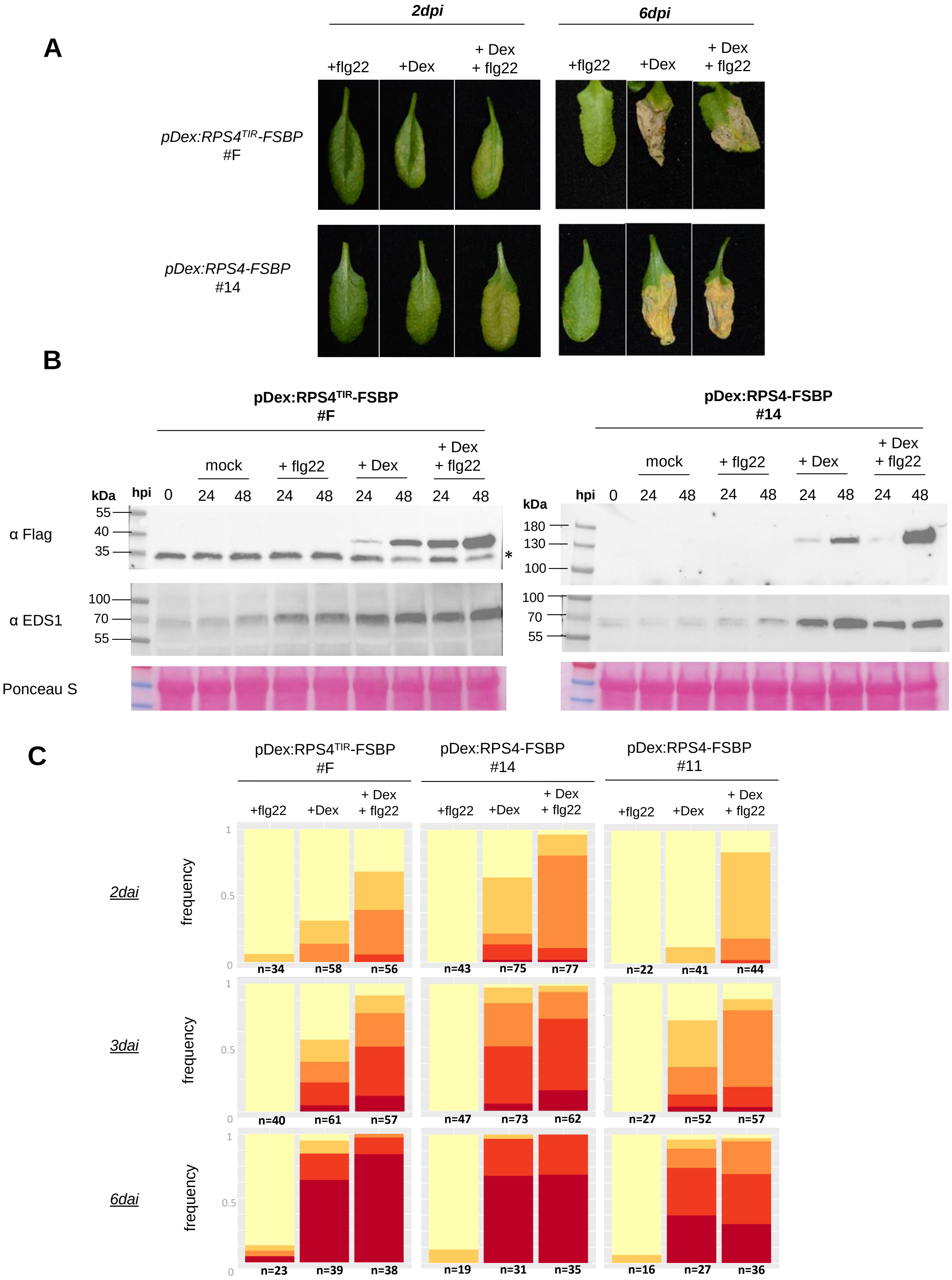
A
2dpi
6dpi
+flg22
+Dex
+ Dex
+ flg22
pDex:RPS4TIR-FSBP
#F
pDex:RPS4-FSBP
#14
B
pDex:RPS4TIR-FSBP
#F
pDex:RPS4-FSBP
#14
mock
+ flg22
+ Dex
+ Dex
+ flg22
kDa
hpi
0
24
48
α Flag
α EDS1
Ponceau S
55
40
35
100
70
180
130
*
C
pDex:RPS4TIR-FSBP
#F
pDex:RPS4-FSBP
#14
pDex:RPS4-FSBP
#11
2dai
3dai
6dai
frequency
1
0.5
0
n=34
n=58
n=56
n=43
n=75
n=77
n=22
n=41
n=44
n=40
n=61
n=57
n=47
n=73
n=62
n=27
n=52
n=57
n=23
n=39
n=38
n=19
n=31
n=35
n=16
n=27
n=36

**Figure S11. Effect of flg22 co-treatment on Dex-induced cell death in Arabidopsis lines carrying *pDex:RPS4$^{TIR}$-FSBP* and *pDex:RPS4-FSBP*.** A. Representative cell death phenotype observed in three week-old Arabidopsis leaves expressing pDex:RPS4$^{TIR}$ or pDex:RPS4, 2 and 6 days after infiltration (dpi) with 100 nM flg22, 20µM Dex or Dex + flg22. B. Immunoblot analysis of RPS4$^{TIR}$-FSBP, RPS4-FSBP and EDS1 in indicated lines, 24 or 48 hours after leaf infiltration with a mock solution (DMSO, -Dex), flg22, Dex or Dex+flg22. Total protein load is indicated by Ponceau red staining. Non-specific bands are indicated by an asterisk. (C) Representation of cell death scores as a percentage of each score (color-coded as indicated in Figure S14), in n>16 leaves of three week-old Arabidopsis lines carrying *pDex:RPS4$^{TIR}$* or *pDex:RPS4* (2, 3 or 6 days after infiltration with 100 nM flg22, 20µM Dex, or Dex + flg22 solutions). Numbers of scored leaves per treatment is indicated at the bottom of the for each condition. Data presented in (C) are the result of at least three independent experiments.

### Table S1: List of transgenic lines used in this study

| line identifier | construct name | genetic background | line number | reference |
|---|---|---|---|---|
| | pDex:RPS4TIR-FSBP | Col-0 | #F | Bernoux et al. 2023 |
| | pDex:L6TIR-FSBP | Col-0 | #4 | this study |
| | pDex:L6MHV-myc | Col-0 | #2 | Bernoux et al. 2023 |
| AT241 | pDex:SNC1TIR-FSBP | Col-0 | #7 | this study |
| | | | #8 | this study |
| AT243 | pDex:L6MHV-FSBP | Col-0 | #18 | this study |
| AT243b | pDex:L6MHV-FSBP | Col-0 | #11 | this study |
| AT244 | pDex:RPS4-FSBP | Col-0 | #11 | this study |
| | | | #14 | this study |
| | | | #22 | this study |
| AT251 | pDex:L6TIR-FSBP | *adr1 triple* (Col-0) | #3 | this study |
| | | | #4 | this study |
| AT252 | pDex:SNC1TIR-FSBP | *adr1 triple* (Col-0) | #1 | this study |
| AT277 | pDex:L6MHV-FSBP | *nrg1 double* (Col-0) | #2 | this study |
| | | | #9 | this study |
| AT278 | pDex:L6MHV-FSBP | *adr1 triple* (Col-0) | #3 | this study |
| AT290 | pDex:L6TIR-FSBP | *adr1 triple* (Col-0) | #4 | this study |
| AT291 | pDex:SNC1TIR-FSBP | *adr1 triple* (Col-0) | #D | this study |
| AT293 | pDex:SNC1TIR-FSBP | *helperless* (Col-0) | #A | this study |
| | | | #2 | this study |
| | | | #3 | this study |
| AT294 | pDex:GFP-MiniTurbo-Flag | Col-0 | #17 | this study |
| AT299 | pDex:L6TIR-FSBP | *nrg1 double* (Col-0) | #A | this study |
| AT300 | pDex:SNC1TIR-FSBP | *nrg1 double* (Col-0) | #C | this study |
| | | | #E | this study |
| | | | #G | this study |
| AT301 | pDex:L6MHV-FSBP | *helperless* (Col-0) | #5 | this study |
| | | | #9 | this study |
| AT309 | pDex:L6MHV-FSBP | *adr1 triple* (Col-0) | #3 | this study |
| AT312 | pDex:L6TIR-FSBP | *helperless* (Col-0) | #4 | this study |
| | | | #5 | this study |
| AT313 | pDex:L6TIR-FSBP | *nrg1 double* (Col-0) | #3 | this study |
| AT330 | pDex:RPS4TIR-FSBP | *helperless* (Col-0) | #5 | this study |
| | | | #6 | this study |
| AT331 | pDex:RPS4TIR-FSBP | *nrg1 double* (Col-0) | #2 | this study |
| | | | #5 | this study |
| | | | #7 | this study |
| | | | #8 | this study |
| AT332 | pDex:RPS4TIR-FSBP | *adr1 triple* (Col-0) | #4 | this study |
| | | | #5 | this study |
| | | | #6 | this study |
| | | | #7 | this study |
| AT336 | pDex:TN2-YFP | Col-0 | #3 | this study |
| | | | #5 | this study |
| AT337 | pDex:RBA1-YFP | Col-0 | #A | this study |
| | | | #2 | this study |
| | pADR1:L2:ADR1 DV-L2-HA | *adr1-l2* (Col-0) | | Roberts et al., 2013 |

### Table S2: List of plasmid constructs used in this study

| Type of plasmid/constructs | Plasmid/construct Name | Insert or PCR product | Primers | Template | Vector Background | Cloning method | Reference |
|---|---|---|---|---|---|---|---|
| Golden gate fragment | MiniTurbo-3flag-35sterm | MiniTurbo-3flag-35sterm | | BsaI compatible fragments were kindly provided by Dr Mbengue (Toulouse University) | | golden gate | This study |
| PCR products | AttB1-RPS4-AttB2 | RPS4 full-length genomic | AttB1-RPS4/AttB2-RPS4 | Arabidopsis Col-0 genomic DNA | | PCR amplification | This study |
| | AttB1-RBA1-AttB2 | RBA1 full-length genomic | MB104/MB105 | Arabidopsis Col-0 genomic DNA | | PCR amplification | This study |
| | AttB1-TN2-AttB2 | TN2 full-length genomic | MB102/MB103 | Arabidopsis Col-0 genomic DNA | | PCR amplification | This study |
| | Kpn1/YFPv-35sterm/Pme1 | Kpn1/YFPv-35sterm/Pme1 | C284/MB29 | pAM35sGWY-YFPv | | PCR amplification | This study |
| | Kpn1/MiniTurbo-3flag-35sterm/Pme1 | Kpn1/MiniTurbo-3flag-35sterm/Pme1 | MB27/MB29 | golden gate fragment MiniTurbo-3flag-35sterm | | PCR amplification | This study |
| GWY vectors | pDex:GWY-3myc | | | | pOpOff2 GWY (Hyg) | | Bernoux et al. 2023 |
| | pDex:GWY-FSBP | | | | pOpOff2 GWY (Hyg) | | Bernoux et al. 2023 |
| | pDex:GWY-YFPv | Kpn1/YFPv-35sterm/Pme1 | | | pOpOff2 GWY (Kan) | restriction ligation | This study |
| | pDex:GWY- | Kpn1/MiniTurbo-3flag-35sterm/Pme1 | | | pOpOff2 GWY (Kan) | restriction ligation | This study |
| pENTRY | pDONR L6 TIR | L6 TIR 1-233 genomic | | | | | Bernoux et al. 2011 |
| | pDONR SNC1 TIR | SNC1 TIR 1-226 cDNA | | | | | Zhang et al. 2017 |
| | pDONR RPS4 TIR | RPS4 TIR 1-235 cDNA | | | | | Williams et al. 2014 |
| | pDONR L6MHV | L6 MHD/V full length genomic | | | | | Bernoux et al. 2016 |
| | pDONR RPS4 | RPS4 genomic sequence (Col-0) | | AttB1-RPS4-AttB2 | pDONR207 | BP Gateway | This study |
| | pDONR207 RBA1 | RBA1 full-length genomic | | AttB1-RBA1-AttB2 | pDONR207 | BP Gateway | This study |
| | pDONR207 TN2 | TN2 full-length genomic | | AttB1-TN2-AttB2 | pDONR207 | BP Gateway | This study |
| Destination vectors for in planta expression | pDex:L6MHV-3myc | L6 MHD/V full-length genomic | | | pDex:GWY-3myc | | Bernoux et al. 2023 |
| | pDex:RPS4TIR-FSBP | RPS4 TIR 1-235 cDNA | | | pDex:GWY-FSBP | | Bernoux et al. 2023 |
| | pDex:L6TIR-FSBP | L6 1-233 genomic | | pDONR L6TIR | pDex:GWY-FSBP | LR Gateway | This study |
| | pDex:SNC1TIR-FSBP | SNC1 TIR 1-226 cDNA | | pDONR SNC1 1-226 | pDex:GWY-FSBP | LR Gateway | This study |
| | pDex:L6MHV-FSBP | L6 MHD/V full-length genomic | | pDONR L6MHV | pDex:GWY-FSBP | LR Gateway | This study |
| | pDex:RPS4-FSBP | RPS4 FL genomic (Col-0) | | pDONR RPS4 | pDex:GWY-FSBP | LR Gateway | This study |
| | pDex:RBA1-YFP | RBA1 full-length genomic | | pDONR207 RBA1 | pDex:GWY-YFP | LR Gateway | This study |
| | pDex:TN2-YFP | TN2 full-length genomic | | pDONR207 TN2 | pDex:GWY-YFP | LR Gateway | This study |

### Table S3: List of primers used in this study

| Gene ID | Targeted gene/construct name | oligo name_ref | sequence (5'-3') | purpose | reference |
|---|---|---|---|---|---|
| AT1G13320 | *PP2AA3* | At1G13320-F qpcr | GACCGGAGCCAACTAGGAC | RT-qPCR | this study |
| | | At1G13320-R qpcr | AAAACTTGGTAACTTTTCC | | |
| AT5G15710 | *galactose oxydase/kelch repeat superfamily protein* | HK10 | TTTCGGCTGAGAGGTTCG | RT-qPCR | this study |
| | | HK10 | GATTCCAAGACGTAAAGCAGA | | |
| AT5G26920 | *CBP60g* | CBP60g-F | TCGTGGACGCCACCACAAACA | RT-qPCR | Kim et al. 2017 |
| | | CBP60g-R | TCAGCGTTCAGCGGCACGAG | | |
| AT1G74710 | *ICS1* | ICS1-R | TTCTGGGCTCAAACACTAAAAC | RT-qPCR | this study |
| | | ICS1-F | GGCGTCTTGAAATCTCCATC | | |
| AT3G48090 | *EDS1* | AtEDS1-F | AGATTATTCAGGTGATCGAGCA | RT-qPCR | Bhandari et al. 2019 |
| | | AtEDS1-R | TTTATGGGCTTGACACTTTGG | | |
| AT2G35980 | *NHL10* | NHL10-qPCR-F | TTCCTGTCCGTAACCCAAAC | RT-qPCR | Boudsocq et al. 2010 |
| | | NHL10-qPCR-R | CCCTCGTAGTAGGCATGAGC | | |
| AT2G19190 | *FRK1* | AtFRK1-F | TGGATCCATCGGTTACCTTG | RT-qPCR | Ngou et al., 2021 |
| | | AtFRK1-R | AGCTTGCAATAGCAGGTTGG | | |
| transgene | *pDex:L6MHV-FSBP, pDex:L6MHV-myc, pDex:L6TIR-FSBP* | qPCR2-L6-F | GCTACTGCTGTTGCCTTGC | RT-qPCR | this study |
| | | qPCR2-L6-R | ACCAGAGGGATTTGTGGAGT | | |
| transgene | *pDex:RPS4-FSBP, pDex:L6MHV-FSBP* | prePoP6prom | ACAACTACAGCTAGCAAGCTTGTCGA | genotyping | this study |
| | | SBPrev | CTAGTCTAAGGCTCTCTTTGTCCT | | |
| transgene | *pDex:L6MHV-FSBP* | L6ex4-5 | GTCGTTGGGGAGACTACCA | genotyping | this study |
| | | SBPrev | CTAGTCTAAGGCTCTCTTTGTCCT | | |
| At1g33560 | *ADR1* | FEK 833 | CAAAGGACGATGATGTTCGAG | genotyping | Saile et al. 2020 |
| | | FEK 834 | CGGATTGTTCACTATAGTAAGG | | |
| At4g33300 | *ADR1 L1* | FEK 836 | ATGGCCATCACCGATTTTTTC | genotyping | Saile et al. 2020 |
| | | FEK 837 | GTCAGGAACAGGATTTCCAG | | |
| At5g04720 | *ADR1 L2* | FEK 829 | tgggagattgtgacacagtc | genotyping | Saile et al. 2020 |
| | | FEK 830 | ATGGCAGATATAATCGGCGG | | |
| At1g33560 | *adr1-1 (T-DNA insertion)* | FEK 834 | CGGATTGTTCACTATAGTAAGG | genotyping | Saile et al. 2020 |
| | | FEK 835 | TTTCATAACCAATCTCGATACAC | | |
| At4g33300 | *adr1-L1-1 (T-DNA insertion)* | FEK 837 | GTCAGGAACAGGATTTCCAG | genotyping | Saile et al. 2020 |
| | | FEK 835 | TTTCATAACCAATCTCGATACAC | | |
| At5g04720 | *adr1-L2-4 (T-DNA insertion)* | FEK 839 | CAACATCTCCTTCACCTTCC | genotyping | Saile et al. 2020 |
| | | VB56 | ATTTTGCCGATTTCGGAAC | | |
| At5g66910 | *nrg1.2* | FEK1068 | CTGGTCTGTATTTTTGGTCCTC | genotyping | Saile et al. 2020 |
| | | FEK1069 | GAAAGTTGTCCTCTCGTATCT | | |
| At5g45250 | *RPS4* | AttB1-RPS4 | GGGGACAAGTTTGTACAAAAAAGCAGGCTTAATGGAGACATCATCTATTTCC | cloning | this study |
| | | AttB2-RPS4 | GGGGACCACTTTGTACAAGAAAGCTGGGTCGAAATTCTTAACCGTGTGCATG | cloning | |
| At1G47370 | *RBA1* | MB104 | CAAAAAAGCAGGCTTAATGacgagcgtgtctcctcggtaca | cloning | this study |
| | | MB105 | CAAGAAAGCTGGGTCaatccttacagtcctgtcatacgtgt | cloning | |
| At1g17615 | *TN2* | MB102 | CAAAAAAGCAGGCTTAATGtattcatcatcgtcttcttcttca | cloning | this study |
| | | MB103 | CAAGAAAGCTGGGTCagaagattcagtcccggatataggt | cloning | |
| | *Kpn1/YFPv-35sterm/Pme1* | C284 | ccccggtaccatggtgagcaagggcgaggagct | cloning | this study |
| | | MB29 | ccccgtttaaacATCTGGATTTTAGTACT | cloning | |
| | *Kpn1/MiniTurbo-3flag-35sterm/Pme1* | MB27 | ccccggtaccATGATTCCATTACTCAACGCA | cloning | this study |
| | | MB29 | ccccgtttaaacATCTGGATTTTAGTACT | cloning | |

**References associated to supplementary data**


Bernoux, M., J. Chen, X. Zhang, K. Newell, J. Hu, L. Deslandes, and P. Dodds. 2023. "Subcellular Localization Requirements and Specificities for Plant Immune Receptor Toll-Interleukin-1 Receptor Signaling." Plant J 114, no. 6 (Jun): 1319-1337. https://dx.doi.org/10.1111/tpj.16195.

Bernoux, M., T. Ve, S. Williams, C. Warren, D. Hatters, E. Valkov, X. Zhang, J. G. Ellis, B. Kobe, and P. N. Dodds. 2011. "Structural and Functional Analysis of a Plant Resistance Protein Tir Domain Reveals Interfaces for Self-Association, Signaling, and Autoregulation." Cell Host Microbe 9, no. 3 (Mar 17): 200-11. https://dx.doi.org/10.1016/j.chom.2011.02.009.

Bhandari, D. D., D. Lapin, B. Kracher, P. von Born, J. Bautor, K. Niefind, and J. E. Parker. 2019. "An Eds1 Heterodimer Signalling Surface Enforces Timely Reprogramming of Immunity Genes in Arabidopsis." Nat Commun 10, no. 1 (02 15): 772. https://dx.doi.org/10.1038/s41467-019-08783-0.

Boudsocq, M., M. R. Willmann, M. McCormack, H. Lee, L. Shan, P. He, J. Bush, S. H. Cheng, and J. Sheen. 2010. "Differential Innate Immune Signalling Via Ca(2+) Sensor Protein Kinases." Nature 464, no. 7287 (Mar 18): 418-22. https://dx.doi.org/10.1038/nature08794.

Kim, J. H., C. D. M. Castroverde, S. Huang, C. Li, R. Hilleary, A. Seroka, R. Sohrabi, D. Medina-Yerena, B. Huot, J. Wang, K. Nomura, S. K. Marr, M. C. Wildermuth, T. Chen, J. D. MacMicking, and S. Y. He. 2022. "Increasing the Resilience of Plant Immunity to a Warming Climate." Nature 607, no. 7918 (Jul): 339-344. https://dx.doi.org/10.1038/s41586-022-04902-y.

Ngou, B. P. M., H. K. Ahn, P. Ding, and J. D. G. Jones. 2021. "Mutual Potentiation of Plant Immunity by Cell-Surface and Intracellular Receptors." Nature 592, no. 7852 (04): 110-115. https://dx.doi.org/10.1038/s41586-021-03315-7.

Roberts, M., S. Tang, A. Stallmann, J. L. Dangl, and V. Bonardi. 2013. "Genetic Requirements for Signaling from an Autoactive Plant Nb-Lrr Intracellular Innate Immune Receptor." PLoS Genet 9, no. 4: e1003465. https://dx.doi.org/10.1371/journal.pgen.1003465.

Saile, S. C., F. M. Ackermann, S. Sunil, J. Keicher, A. Bayless, V. Bonardi, L. Wan, M. Doumane, E. Stöbbe, Y. Jaillais, M. C. Caillaud, J. L. Dangl, M. T. Nishimura, C. Oecking, and F. El Kasmi. 2021. "Arabidopsis Adr1 Helper Nlr Immune Receptors Localize and Function at the Plasma Membrane in a Phospholipid Dependent Manner." New Phytol 232, no. 6 (Dec): 2440-2456. https://dx.doi.org/10.1111/nph.17788.

Saile, S. C., P. Jacob, B. Castel, L. M. Jubic, I. Salas-Gonzáles, M. Bäcker, J. D. G. Jones, J. L. Dangl, and F. El Kasmi. 2020. "Two Unequally Redundant "Helper" Immune Receptor Families Mediate Arabidopsis Thaliana Intracellular "Sensor" immune Receptor Functions." PLoS Biol 18, no. 9 (09): e3000783. https://dx.doi.org/10.1371/journal.pbio.3000783.

Williams, S. J., K. H. Sohn, L. Wan, M. Bernoux, P. F. Sarris, C. Segonzac, T. Ve, Y. Ma, S. B. Saucet, D. J. Ericsson, L. W. Casey, T. Lonhienne, D. J. Winzor, X. Zhang, A. Coerdt, J. E. Parker, P. N. Dodds, B. Kobe, and J. D. Jones. 2014. "Structural Basis for Assembly and Function of a Heterodimeric Plant Immune Receptor." Science 344, no. 6181 (Apr 18): 299-303. https://dx.doi.org/10.1126/science.1247357.

Zhang, X., M. Bernoux, A. R. Bentham, T. E. Newman, T. Ve, L. W. Casey, T. M. Raaymakers, J. Hu, T. I. Croll, K. J. Schreiber, B. J. Staskawicz, P. A. Anderson, K. H. Sohn, S. J. Williams, P. N. Dodds, and B. Kobe. 2017. "Multiple Functional Self-Association Interfaces in Plant Tir Domains." Proc Natl Acad Sci U S A 114, no. 10 (Mar): E2046-E2052. https://dx.doi.org/10.1073/pnas.1621248114.
